# Interconnected assembly factors regulate the biogenesis of mitoribosomal large subunit

**DOI:** 10.1101/2020.06.28.176446

**Authors:** Victor Tobiasson, Ondřej Gahura, Shintaro Aibara, Rozbeh Baradaran, Alena Zíková, Alexey Amunts

## Abstract

Mitoribosomes consist of ribosomal RNA and protein components, coordinated assembly of which is critical for function. We used mitoribosomes with reduced RNA and increased protein mass from *Trypanosoma brucei*, to provide insights into the biogenesis of mitoribosomal large subunit. Structural characterisation of a stable assembly intermediate revealed 22 assembly factors, some of which are also encoded in mammalian genomes. The assembly factors form a protein network that spans over 180 Å, shielding the ribosomal RNA surface. The entire central protuberance and L7/L12 stalk are not assembled, and require removal of the factors and remodeling of the mitoribosomal proteins to become functional. The conserved proteins GTPBP7 and mt-EngA are bound together at the subunit interface in proximity to the peptidyl transferase center. A mitochondrial acyl-carrier protein plays a role in docking the L1 stalk which needs to be repositioned during maturation. Additional enzymatically deactivated factors scaffold the assembly, while the exit tunnel is blocked. Together, the extensive network of the factors stabilizes the immature sites and connects the functionally important regions of the mitoribosomal large subunit.

## Introduction

Mitoribosomes differ from bacterial and cytosolic ribosomes in their ribosomal RNA (rRNA), protein content, overall size, and structure. Their formation is an intricate and hierarchical process involving multiple proteins and RNA molecules working in coordination and under tight regulation (Pearce et al 2017). The cooperative effort involves regulation of two genomes, because rRNA is encoded by the organellar genome, and almost all the mitoribosomal proteins and assembly factors are encoded by the nuclear genome and therefore imported from the cytosol (Couvillion et al 2016). Finally, the fundamental process of the mitoribosomal assembly is complicated due to the localization of its large subunit (mtLSU) to the inner mitochondrial membrane. Therefore, stages of assembly were suggested to involve specific steps and kinetics (Bogenhagen et al 2014; Antonicka and Shoubridge 2015; De Silva et al 2015). The presence of different compositions is hypothesized to promote formation of defined pre-mitoribosomal complexes with as-yet-unknown organelle-specific auxiliary factors.

Mitochondria of *Trypanosoma brucei* provide a good model for studying the assembly process, because their mitoribosomes consist of over a hundred components, and the ratio of protein to rRNA is unusually high (Zikova et al 2008; Ramrath et al 2018). Since the rRNA forms a compact core of the mitoribosome, and proteins are mostly peripherally associated, an architecture based on the reduced rRNA and supernumerary mitoribosomal proteins would need additional stabilization for its assembly. Therefore, it increases the chances to characterize defined pre-mitoribosomal complexes, which are not stable enough for biochemical isolation in mitochondria of other species. Indeed, structural characterization of an assembly intermediate of the *T. brucei* mitoribosomal small subunit (mtSSU) provided insight into its assembly pathway with many newly detected proteins (Saurer et al 2019).

The mtLSU accommodates the peptidyl transferase center (PTC) that catalyzes formation of peptide bonds between amino acids, tRNA binding sites, the L7/L12 stalk that is responsible for the recruitment of translation factors, the L1 stalk, the central protuberance (CP) that facilitates communication between various functional sites, and the exit tunnel to egress a synthesized protein. In bacteria, our understanding of the LSU assembly is relatively limited (Davis and Williamson 2017). It comes primarily from a characterization of the final maturation stages (Li et al 2013; Jomaa et al 2014; Ni et al 2016), studies on incomplete LSU particles as a result of protein depletion (Davis et al 2016), as well as *in vitro* reconstitution studies with purified ribosomal RNA and protein components (Nikolay et al 2018). These studies identified different LSU precursors with assembly factors bound to rRNA components (Davis and Williamson 2017). In mitochondria, the mtLSU lacks many of the rRNA components involved in the canonical pathways, and higher complexity of the interactions between the mitoribosomal proteins at the functional sites has evolved (Ott et al 2016; Greber and Ban 2016). A functional mtLSU requires a folded rRNA core, a flexible L1 stalk that is involved in tRNA movement, an extended L7/L12 protrusion for binding of translational factors, and a proteinaceous CP formed by mitochondria-specific elements involved in tRNA binding (Aibara et al 2020; Tobiasson and Amunts 2020). However, only the final stage of the mtLSU assembly with fully mature functional sites has been visualized (Brown et al 2017; Itoh et al 2020), and no preceding steps in the formation have been detected. Therefore, mtLSU assembly remains poorly understood.

To provide insight into the process of the mtLSU assembly, we determined the cryo-EM structure of a native *T. brucei* mtLSU assembly intermediate (pre-mtLSU) in a complex with assembly factors. Most of the assembly factors have not been previously implicated in mitoribosomal biogenesis. The structural data suggests that the biogenesis relies on an extensive protein network formed by the assembly factors that connect a premature PTC, the L1 and L7/L12 stalks with the CP, while the exit tunnel is blocked. A homology search suggests that some of the newly identified assembly factors are also conserved in mitochondria from other species, including mammals, and therefore may represent general principles. Comparison with two bacterial assembly intermediates (Zhang et al 2014; Seffouh et al 2019) further provides insights into the conserved GTPases GTPBP7 and mt-EngA bound at the subunit interface.

## Results

### Structural determination and composition of the native pre-mtLSU complex

We used a *T. brucei* procyclic strain grown in low-glucose medium that maintains translationally active mitochondria. Mitoribosomal complexes were purified directly from *T. brucei* mitochondria and analyzed by single-particle cryo-EM. During image processing, in addition to the intact monosomes, we detected a pool of free subunits. We focused the analysis on this population and through 3D classification isolated a homogeneous subset of pre-mtLSUs that corresponded to ~3.5 % of the particles combined from five data sets.

896,263 particles were picked using Warp (Tegunov and Cramer 2019), and further processed using RELION (Kimanius et al 2016; Zivanov et al 2018). We performed reference-based 3D classification with references generated from a preliminary classification of a screening data set. This resulted in 207,788 particles corresponding to the mtLSU shape but distinct from that of a mature mtLSU of which we found 152,816 particles. Refinement of those assigned and subsequent classification using fine-angular searches with a solvent mask identified 32,339 pre-mtLSU particles (Appendix Fig S1). To improve the angles further, the particles were subjected to masked auto-refinement. Following the CTF refinement, we obtained a reconstruction of a pre-mtLSU that likely reflects a stable premature complex. This was evidenced by the presence of densities corresponding to conserved ribosomal assembly factors.

The cryo-EM reconstruction was refined to 3.50 Å resolution (Table S1). This allowed us to build a ~2.2 MDa model and assign assembly factors, as well as additional mitoribosomal proteins, directly from the density (Figs 1 and 2, Appendix S2). Six distinct features define the overall pre-mtLSU; 1) the rRNA domain V is well resolved and covered by newly identified mitochondria-specific assembly factors, 2) the subunit interface consists almost entirely of newly identified assembly factors and two conserved GTPases, 3) the proteinaceous CP is absent, 4) the L7/L12 stalk proteins are missing, and its rRNA platform is not folded, instead assembly factors occupy similar positions, 5) the L1 stalk is shifted inward ~30 Å and linked to the CP base by assembly factors, and 6) the exit tunnel is blocked. Due to these features, compositional and conformational changes are required for the maturation of the pre-mtLSU. In terms of the mitoribosomal proteins, 18 previously identified proteins are missing from the structure of the pre-mtLSU. Seven of these have bacterial homologs (uL10m, uL12m, uL16m, bL27m, bL31m, bL33m and bL36m) and the rest are mitochondria specific (Fig 1, Table S2). Additionally, we assigned sequences to previously unidentified mtLSU proteins uL14m and mL101 (Fig 2).

**Figure 1.**
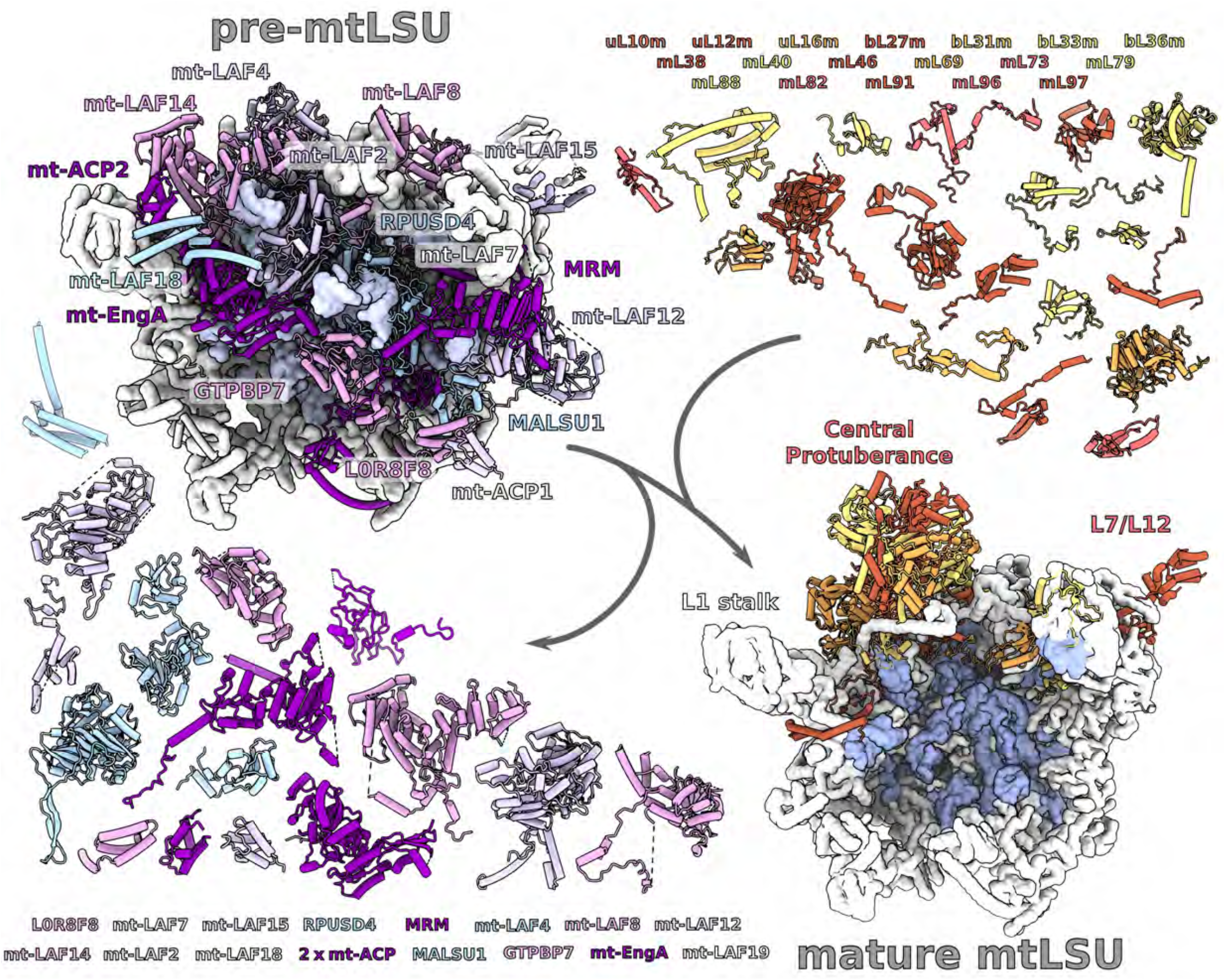
Structure of *T. brucei* pre-mtLSU with assembly factors. Left, the overall modeled structure of the pre-mtLSU (rRNA shown as surface) with models of assembly factors (helical tubes, shades of purple) covering the subunit interface, CP, L7/L12 stalk and connecting to the L1 stalk. Right, structure of the mature mtLSU (PDB ID 6HIX) with 18 additional mitoribosomal proteins (shades of orange) absent from pre-mtLSU.

**Figure 2.**
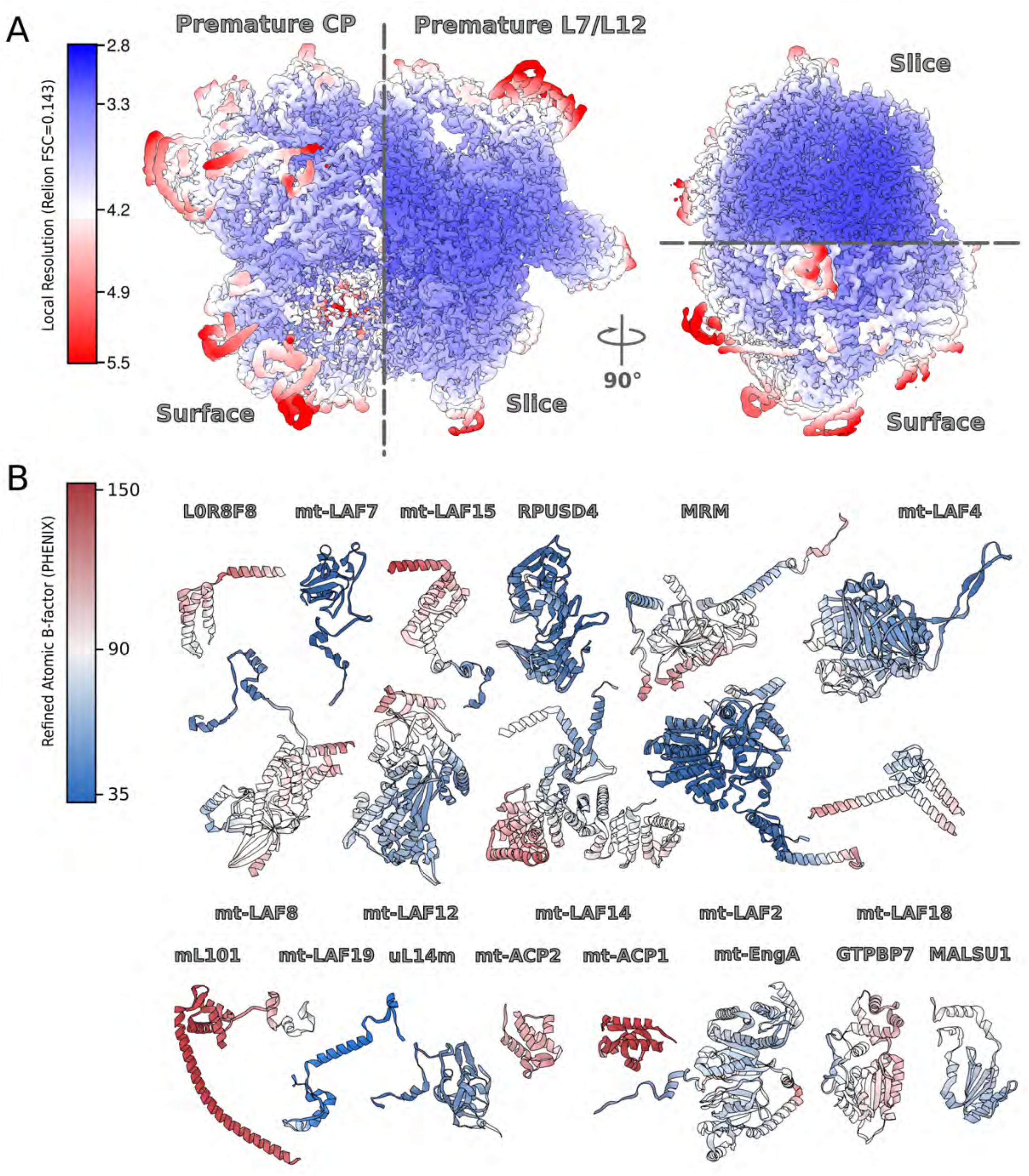
Cryo-EM data quality. (**A**) Final map colored by local resolution. (**B**) Models for individual assembly factors and newly identified proteins colored by refined atomic B-factor.

Following the previously identified mitoribosomal small subunit assembly factors (Saurer et al 2019), we adopt a similar nomenclature for the mitochondria specific large subunit factors. Therefore, we refer to them as mitochondrial Large subunit Assembly Factor(s) (mt-LAF). Proteins with mitochondrial homologs are referred to as their human names. Proteins with bacterial homologs but not identified in humans are referred to as their bacterial names with the prefix “mt-“. The identified assembly factors of the mitoribosome include two homologs of bacterial GTPase assembly factors GTPBP7 (RbgA in bacteria) and mt-EngA, a homolog of the ribosome silencing factor mt-RsfS (MALSU1), a DEAD-box RNA helicase (mt-LAF2), two pseudouridinases, RPUSD4 and mt-LAF4, as well as a methyltransferase MRM, two copies of the mitochondrial acyl-carrier protein mt-ACP, two LYR-motif-containing proteins L0R8F8 and mt-LAF18. Finally, six other proteins with previously unassigned functions mt-LAF7, 8, 12, 14, 15, 19 are present. In the model, we included only the parts for which backbone geometry is apparent. Other regions with only partial or poor density visible were modeled as UNK1-11.

### GTPase mt-RbgA (GTPBP7) is structurally linked to the mitoribosomal core via specific assembly linkers

We started the structural analysis by searching for similar assembly intermediate architectures in bacterial counterparts. Particles with an absent CP were reported previously in RbgA-depleted *Bacillus subtilis* cells. RbgA was then added *in vitro* and shown to bind to the complex, which identified its role as an assembly factor (Seffouh et al 2019). RbgA belongs to the Ras GTPase family typically containing a low intrinsic GTPase activity which is increased in the presence of a mature LSU subunit (Achila et al 2012). It has an N-terminal GTPase domain and a C-terminal helical domain that forms a five-helix bundle (Pausch et al 2018). In the pre-mtLSU structure, we found a conserved mitochondrial homolog of RbgA, GTPBP7. Studies in yeasts reported that deletion of this protein (Mtg1 in yeast) results in respiration deficiency (Barrientos et al 2003). In *B. subtilis*, where this assembly factor is essential, the LSU:RbgA complex showed that the N-terminal domain overlaps with rRNA H69 and H71, and that the C-terminal helical domain interacts with H62 and H64 (Seffouh et al 2019). In this position, RbgA displaces the P-site and further interacts with the surrounding rRNA, including H92 and H93. Therefore, the binding of RbgA requires specific contacts with rRNA. In our map of the *T. brucei* pre-mtLSU, the corresponding rRNA regions forming the binding site for GTPBP7 are not observed. However, the comparison of our structure with the *B. subtilis* LSU:RbgA complex (PDB ID 6PPK) shows nearly identical conformation of the factor on the pre-mtLSU complex (Fig EV1). This includes the peripheral interaction between the GTPBP7 C-terminal domain and the mitoribosomal protein uL14m (Fig 3). In addition, the position of the catalytic GTPase site is also conserved, although the nucleotide binding site of GTPBP7 is empty (Fig 3B). A mutational analysis previously identified His67 (His9 in *B. subtilis*) as a key catalytic residue, and its correct conformation is guided by rRNA (Gulati et al 2013). Despite the overall conservation in mitochondria, the rRNA that is proposed to position the residue in bacteria is missing in our pre-mtLSU structure.

**Figure 3.**
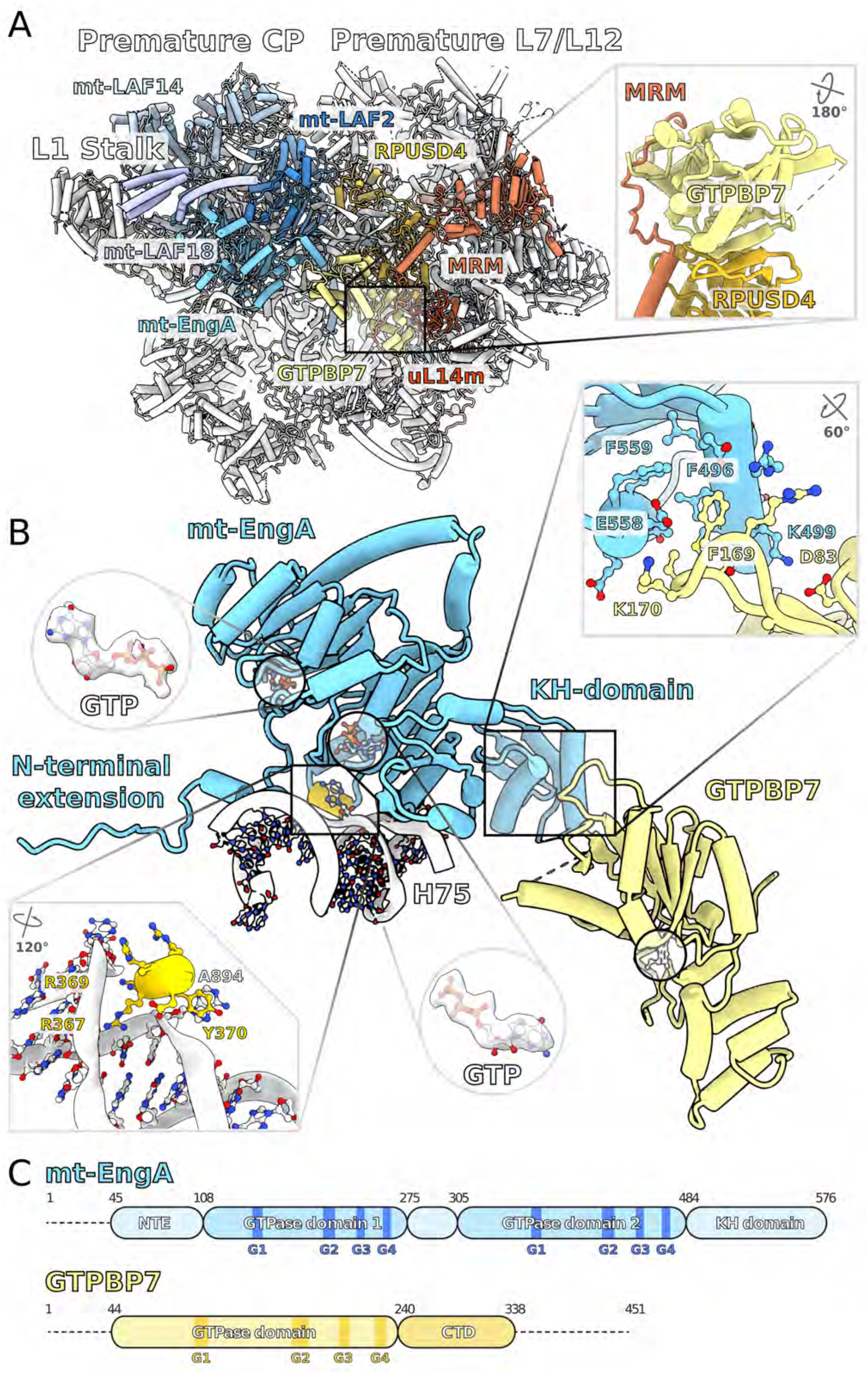
Binding of the GTPBP7 and mt-EngA to the subunit interface. (**A**) GTPBP7 (yellow) is bound to RPUSD4 and MRM, which are connected to the L7/L12 stalk; mt-EngA (blue) is associated with mt-LAF2 and mt-LAF18, which are connected to the CP. (**B**) A short helix of mt-EngA (yellow) interacts with a flipped A894 nucleotide from H75 (white). Two GTPs in their binding sites on mt-EngA are shown as sticks. Absent GTP displayed in its binding site on GTPBP7 is shown as white sticks. The residues forming interactions between mt-EngA and GTPBP7 are shown in the top right in set. (**C**) Schematic representation of mt-EngA and GTPBP7 indicating the positions of the conserved GTP binding motifs.

We found that the conserved position of GTPBP7 in *T. brucei* is maintained through two specialized assembly linkers (Fig 3A). The first linker is established between the C-terminal domain and the MRM N-terminal helix. The latter adopts a crescent shape around the C-terminal domain of GTPBP7, forming a series of contacts with four out of its five helices (Fig 3A). The second linker is provided by RPUSD4 approaching from the mitoribosomal core. It interacts with the GTPBP7 C-terminal domain and contributes a β-strand to a shared β-sheet (Fig EV2B). Therefore, GTPBP7 is anchored to the flexible rRNA core via two dedicated factors that compensate for the lack of rRNA contacts.

RPUSD4 belongs to a family of site-specific RluD pseudouridine synthases involved in the bacterial LSU assembly and responsible for creating of pseudouridines at positions 1911, 1915 and 1917 *(E. coli* numbering) in the H69 end-loop (Gutgsell et al 2001; Gutgsell et al 2005). In our pre-mtLSU structure, RPUSD4 encircles the immature rRNA nucleotides A1008-C1013 as well as U1075-U1086 with the connecting nucleotides being unstructured (Figs EV2A and EV4). Its active site is occupied by cytosine C1010 of H90 forming hydrogen bond with glutamic acid E316 (Fig EV2A), suggesting lack of catalytic activity in the detected state. The N-terminal domain of RPUSD4 is positioned at the distance of ~80 Å facing towards the L7/L12 stalk. Thus, *T. brucei* RPUSD4 performs a stabilizing role for GTPBP7 at the subunit interface and connects with the L7/L12 stalk to coordinate the maturation of the different functional sites (Fig 4A,B).

**Figure 4.**
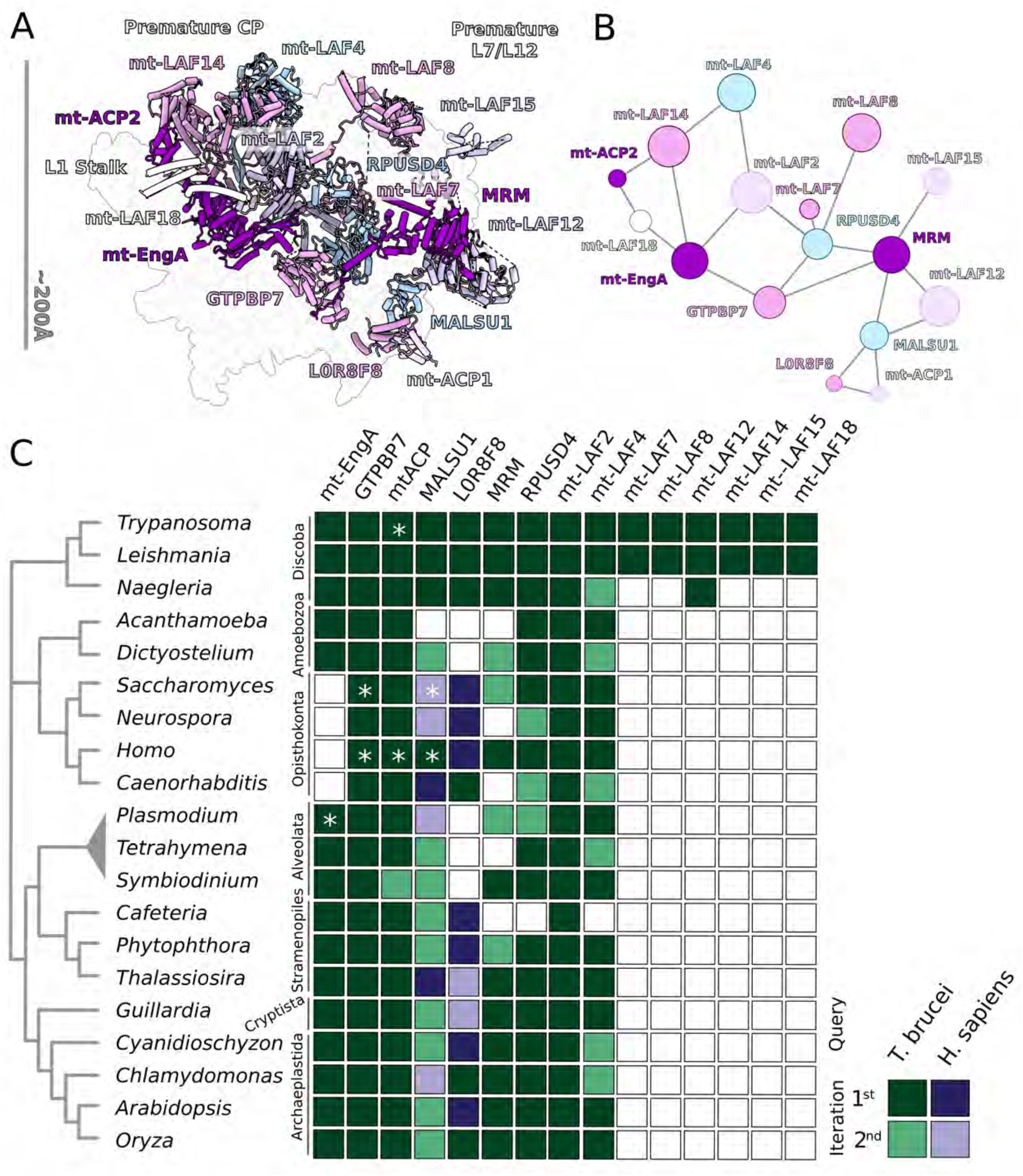
Network of interactions between the assembly factors in pre-mtLSU. (**A**) Assembly factors shown on the background of the pre-mtLSU density map, featuring the interconnection. (**B**) Schematic of protein-protein network. The node size represents the molecular mass of the protein. All the assembly factors are linked in a continuous network. (**C**) Homology search of the assembly factors. Colored squares indicate identified homologs/orthologs using *T. brucei* (green) or human (purple) assembly factors as queries. White squares indicate not-identified homologs/orthologs. The stars mark proteins, for which experimental data has been reported.

MRM belongs to the family of SpoU RNA methyltransferases, but appears to have a closed active site obstructed by Phe334, Arg327 and Glu417 that prevents the binding of the typical Sadenosyl methionine cofactor (Fig EV2B), and the sequence of the conserved motif (Hori 2017) is disrupted (Appendix Fig S3). It is located peripherally, and bound to the mitoribosome via a C-terminal 24-residue helix interacting with rRNA H41/42, and via contacts with mt-LAF12 (Fig EV2B).

Together, RPUSD4 and MRM/mt-LAF12 perform a structural scaffolding role for binding GTPBP7. A homology search of the assembly factors reveals that RPUSD4 and MRM are also present in most eukaryotes (Fig 4C). Since GTPBP7 is present in other organisms, our data suggests the reported cooperative action of the assembly factors might be conserved.

### GTPase mt-EngA is stabilized via protein extensions

In the subunit interface, we identified another conserved GTPase homolog, mt-EngA. It contains two GTPase domains arranged in tandem as well as a C-terminal K homology (KH) domain which is pointed towards the PTC. We could model two GTPs in the GTPase domains (Fig 3B). The overall position of mt-EngA is identical to bacteria, suggesting functional conservation. The assembly factor occupies the space between the PTC and the E-site (Fig EV1), and a role in chaperoning rRNA has been proposed (Zhang et al 2012). However, the comparison with *E. coli* LSU:EngA complex reveals conformational differences that highlight the nature of the mitochondrial protein-rich system, and its role in the stabilization of the conserved assembly factor.

Firstly, the N-terminal GTPase domain is extended by 60 residues, with residues 101-108 stabilizing a helix-turn-helix motif (275-305), which remained unresolved in the bacterial complex (Fig EV1B). The N-terminal extension is generally present in mitochondria from other species (Appendix Fig S4). This motif is important for the stabilization of mt-EngA, because one helix is stacked against a helix of mt-LAF2, whereas the other forms a helical bundle with mt-LAF14 (Fig EV3).

Secondly, the N-terminal residues 72-75 of EngA stabilize a short helix (residues 367-374), which is buried within rRNA groove via Arg367 and Arg369 (Fig 3B). It disrupts the local structure of H75 and stabilizes the flipped nucleotide A894. This loop is also highly charged in the corresponding *E.coli* structure, but does not adopt the helical conformation observed here. Finally, the N-terminus forms additional contacts with five mitoribosomal proteins (bL28m, bL35m, bL19m, mL64, mL74), a stabilizing protein mass that compensates for the missing rRNA in this region. Overall, while the N-terminal GTPase domain aligns well with the bacterial EngA, its interacting partners in our structure are more proteinaceous and specific to mitochondria.

The conserved globular domains of mt-EngA are associated with the pre-mtLSU core via mt-LAF14. Its three helices from the N-terminus encloses the N-terminal GTPase domain helix 230-242 (Fig EV3). Here, mt-LAF14 replaces the missing rRNA H82-87 and protein L1, which binds the EngA N-terminal GTPase domain in bacteria. Factor mt-LAF14 spans over 100 Å to the top of the CP, where it also stabilizes unwound rRNA (Figs 4, EV4). Thus, mt-EngA is bound via a protein extension and also associated with the protein-based scaffold of assembly factors, including the high molecular weight mt-LAF2 and mt-LAF14, which are connected to the CP.

### The module GTPBP7:mt-EngA coordinates maturation of interfacial rRNA

The process of the LSU assembly is dynamic with a cooperative action of different assembly factors (Davis et al 2016; Davis and Williamson 2017). Although GTPBP7 and EngA have previously been visualized separately on the bacterial LSU through deletion and reconstitution experiments, our cryo-EM structure shows both factors simultaneously associated with the pre-mtLSU and with each other. The presence of both factors rationalizes why rRNA domain V is better resolved than in the mature mt-LSU (Fig 5). We were able to model 33% more nucleotides relative to the mature mt-LSU, which shows that the H89-93 region does not occupy the expected bacterial position and highlights a need for prominent remodeling during maturation (Appendix Figs S5 and S6).

**Figure 5.**
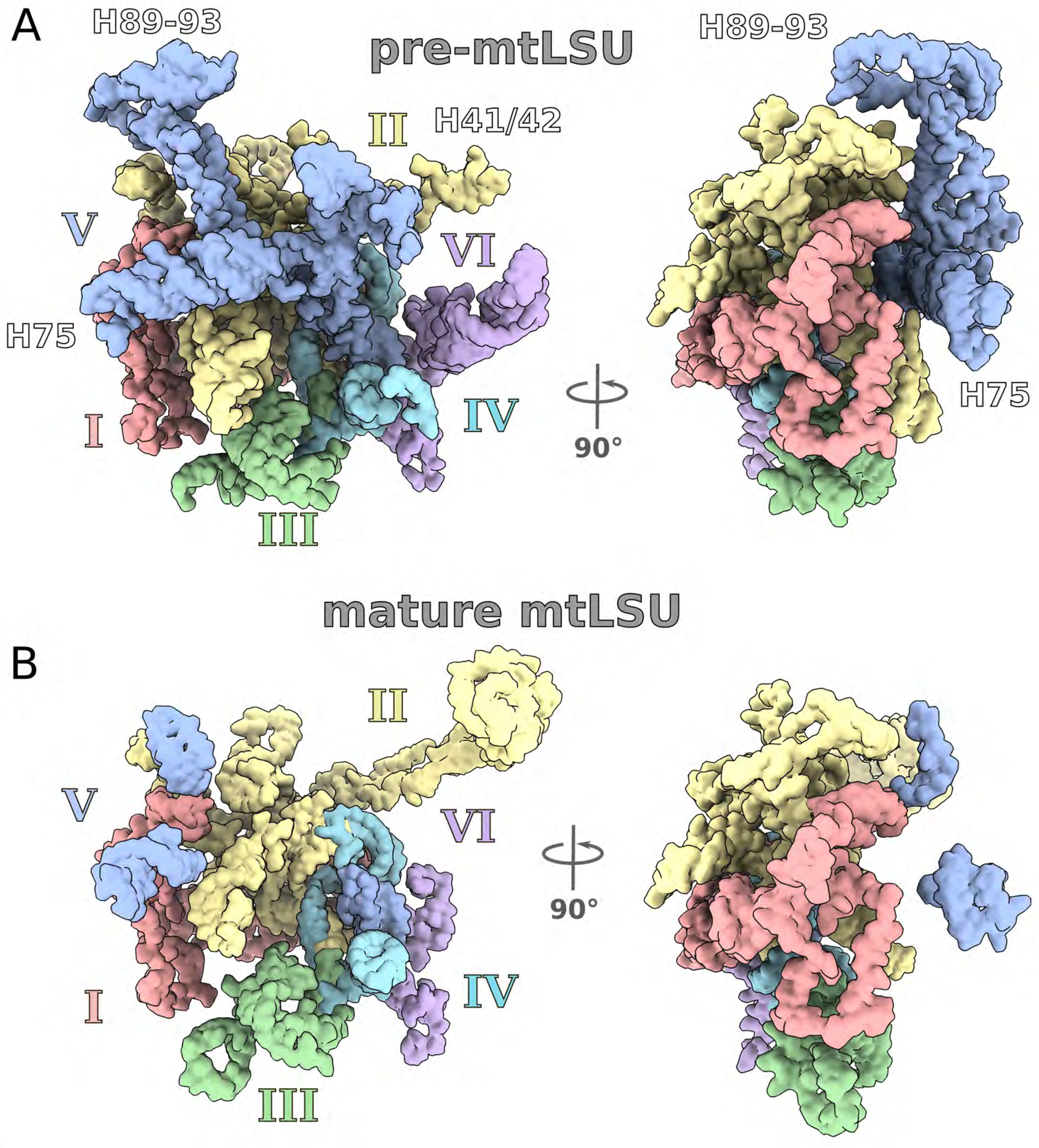
Tertiary structure of rRNA in pre-mtLSU (A) and mature mtLSU (B). Density map lowpass-filtered to 5 Å for clarity shown from the subunit interface (left) and sideview (right). Two views of rRNA related by 90° are shown with each domain in a different color. Domain V is more structured in pre-mtLSU, and H89-93 adopt a different conformation. Domain II that is responsible for L7/L12 stalk is largely disordered.

The contacts between GTPBP7 and mt-EngA are formed via the N-terminal domain and KH domains, respectively (Fig 3B). The shared surface area is ~500 Å, and each of the domains is also associated with rRNA. The contacts formed between GTPBP7 and mt-EngA include electrostatic interactions, as well as hydrophobic residues (Fig 3B). Since the structures and positions of both factors are conserved with bacteria, and we identified homologs in representative eukaryotic species, these results indicate that the simultaneous binding might be a conserved feature.

### DEAD-Box helicase mt-LAF2

In the region connecting the CP with the body of the pre-mtLSU, a conserved ATP-dependent DEAD-box (Asp-Glu-Ala-Asp) RNA helicase was found, namely mt-LAF2. It belongs to a large family of RNA helicases that unwind short RNA duplexes and participate in different aspects of cellular processes, including cell cycle regulation, apoptosis, and the innate immune response (Xing et al 2019).

Factor mt-LAF2 is one of the largest mitoribosomal assembly factors (Fig 2B), spanning 110 Å through the rRNA core to the CP (Figs 4 and 6, Figure EV4). It contains two helicase domains; a DEAD-box and a Helicase C domain (helicase 1 and 2, Fig 6). The two helicases together form the conserved fold for RNA and ATP binding with a linker between them. Typical terminal extensions are also present, and the extended C-terminus anchors mt-LAF2 to the rRNA core by forming contacts at the interface between premature rRNA domains II, IV and V, implying the factor associates early in biogenesis.

**Figure 6.**
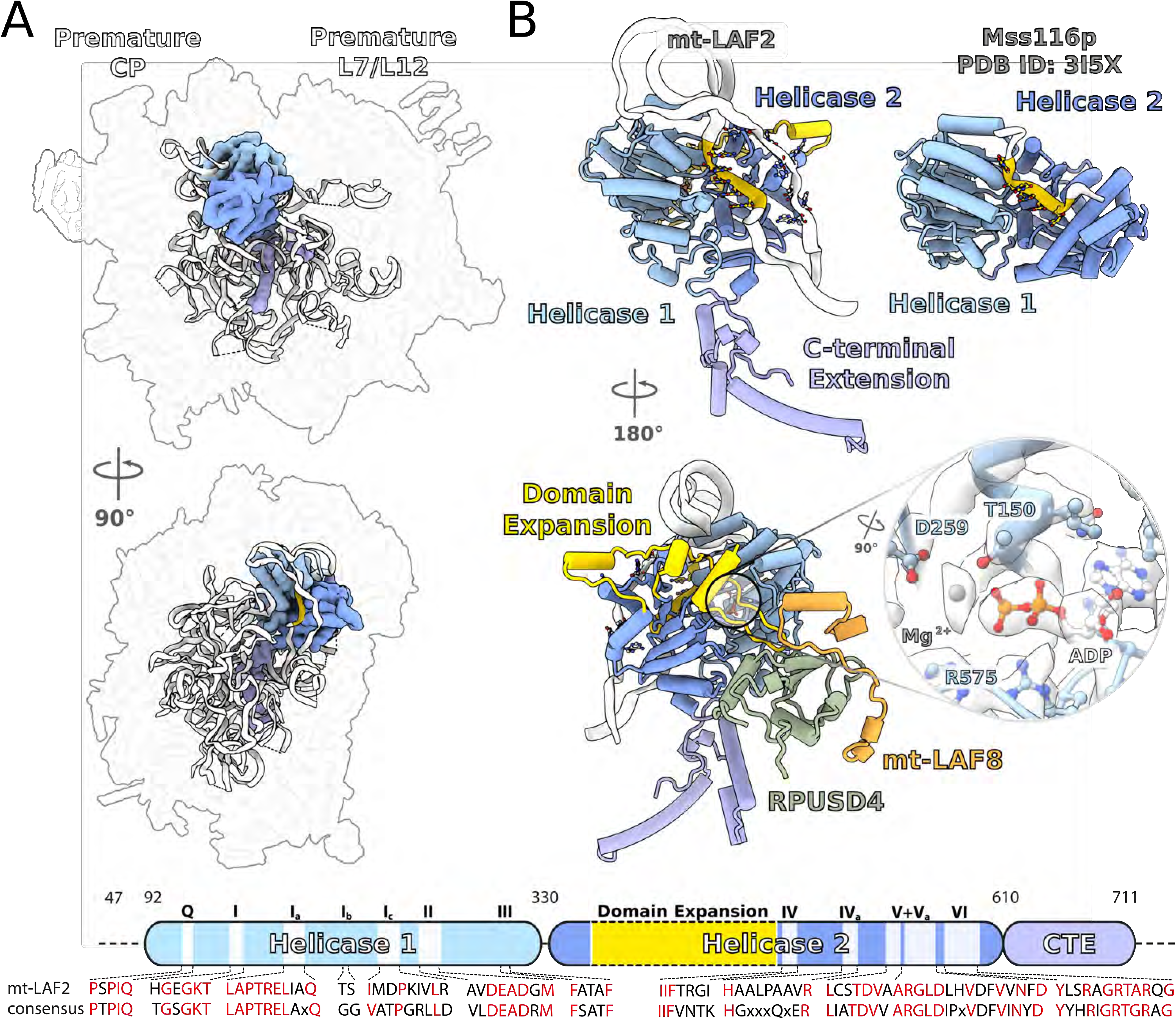
DEAD-Box helicase mt-LAF2 is buried in the pre-mtLSU in closed conformation with bound ADP. **(A)** Relative placement of mt-LAF2 (surface) bound to rRNA (white ribbon). Helicase domain 1 (DEAD box) is light blue, helicase domain 2 (Helicase C) is blue, terminal extensions are purple. **(B)** Comparison with yeast Mss116p shows that rRNA is bound to mt-LAF2 via its phosphate backbone in a similar mode (yellow). Helicase domain 2 expansion in mt-LAF2 (yellow) shields the ADP, and stabilized by RPUSD4 and mt-LAF8. The density in the binding pocket (inset) corresponds to ADP and Mg^2+^ ion. Schematic representation of mt-LAF2 indicating conserved regions is shown in the bottom panel.

Comparison with yeast Mss116p (Del Campo et al 2009) reveals that in contrast to the archetypal DEAD-box RNA helicase, mt-LAF2 has an 120 residue expansion in the Helicase domain 2 that shields the nucleotide moiety (Fig 6B). The helicase core is in a closed conformation, tightly binding the rRNA H80 region. The rRNA is bound via its phosphate backbone, similarly to Mss116p. In the mature human mtLSU, this region forms the P-loop, a constituent of the peptidyl-tRNA binding site (Aibara et al., 2020). Therefore, mt-LAF2 prevents the formation of the functional site for tRNA binding in mtLSU.

The adenosine nucleotide is well resolved in the map, and the density in the binding pocket corresponds to adenosine diphosphate (ADP) (Fig 6B), whereas no continuous density for γ-phosphate is found. The ADP is coordinated by the residues Thr150, Asp295, Arg575, and an Mg^2+^ ion (Fig 6B). ATP hydrolysis was shown to promote substrate release and trigger dissociation and regeneration of the enzyme (Liu et al 2008; Henn et al 2010). However, in our structure, the helicase 2 expansion forms a tertiary junction with two α-β folds from the deactivated RPUSD4 and mt-LAF8 (Fig 6B). This architecture would interfere with the release of the ADP and substrate from mt-LAF2. The interactions further prevent mt-LAF2 disassociation from the pre-mtLSU in the observed state. Therefore, RPUSD4 and mt-LAF8 play a structural role in stabilizing the assembly intermediate.

### Maturation of the L7/L12 stalk

The L7/L12 stalk is a universal mobile element that associates translational protein factors with the A-site. It typically consists of the rRNA platform that connects to the flexible factor-recruiting series of bL12m copies. The ubiquitous ribosomal proteins uL10m, uL11m and mitochondria-specific mL43 link the different components together. In our structure, most of the protein constituents of the stalk are missing and others adopted conformational changes (Fig 7A).

**Figure 7.**
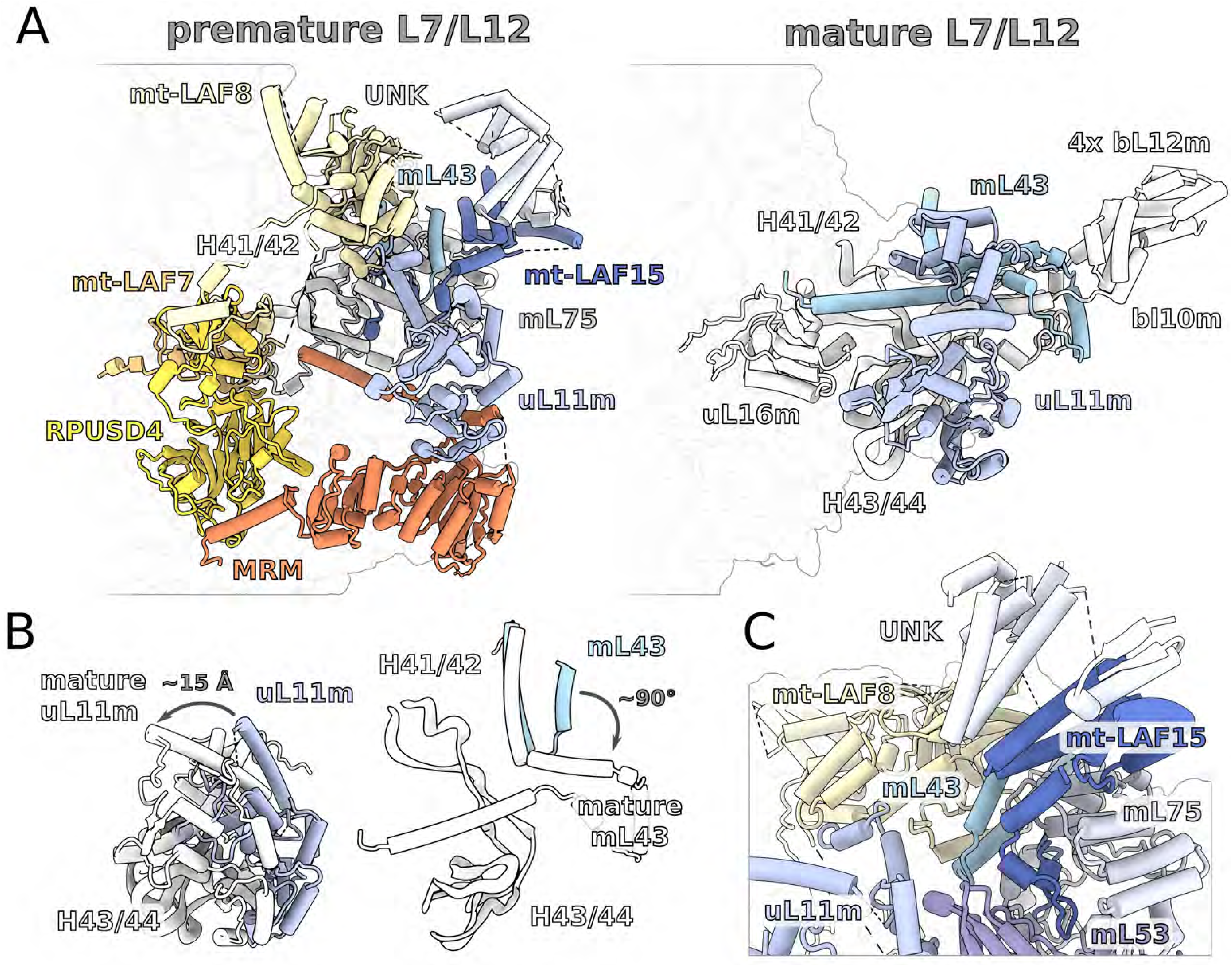
Assembly of the L7/L12 stalk. (**A**) In pre-mtLSU, RPUSD4 extends from the subunit interface to occupy the position of uL16m in the mature mtLSU. Factors mt-LAF7 and mt-LAF8 are bound at the stalk base to the unfolded rRNA H41/42. Factor mt-LAF15 and an additional protein UNK6 form a protrusion similar to bL10m:bL12m. Other mitoribosomal proteins removed for clarity. (**B**) Conformational changes from pre-mtLSU (blue) to mature mtLSU (white) include mL43 and uL11m. (**C**) mt-LAF15, mL75, and UNK6 protein form continuum of at least 13 helices that is peripherally associated.

In the region of the platform, at least three proteins (uL16m, bL36m, mL88) associated with the rRNA in the mature mtLSU are absent. Consistently, the rRNA platform is not folded, as the folding relies on the missing mitoribosomal proteins. Instead, the N-terminal domain of RPUSD4 extends from the subunit interface to occupy part of the space left by uL16m absence (Fig 7A and B). It binds two specific assembly factors mt-LAF7 and mt-LAF8. Factor mt-LAF8 mediates further contacts with the core of the pre-mtLSU. It consists of 7-stranded beta-barrel, 12 α-helices, and 63-residue tail inserted into the mitoribosomal core. Our pre-mtLSU structure suggests that both mt-LAF7 and mt-LAF8 need to dissociate for the missing mitoribosomal proteins to assemble (Fig 7A).

In the factor-recruiting region, instead of the terminal bL12m copies, mt-LAF15 and density corresponding to UNK6 form a protrusion (Fig 7A). They comprise a protein continuum of at least 13 helices associated with each other. This assembly is attached to the platform region through a 30-residue C-terminal tail of mt-LAF15, forming a helical bundle with mL75 (Figs 7A and 7C). Overall, this protein module extends ~70 Å from the core in a similar fashion to the L7/L12 stalk, but appears to be mutually exclusive.

The position of the uL10m N-terminus, which links the stalk to the body in the mature mt-LSU, is occupied by mt-LAF15 C-terminus. It interacts with mL43, resulting in its helix being bent by 90° (Fig 7B). This conformational change and the lack of uL10m is correlated with ~15 Å shift of uL11m to occupy the formed void (Fig 7B). Nevertheless, mt-LAF15 is only peripherally associated with mL43, and it cannot be excluded that the protrusion is independently replaced by the mature L7/L12 stalk.

Based on the structural information, the following working model for the L7/L12 stalk maturation, which includes dismantling, remodeling and assembly can be proposed (Fig EV5): 1) RPUSD4, which is extended from the subunit interface anchoring GTPBP7, has to be removed, 2) mt-LAF7:mt-LAF8 has to be released from the ribosomal core, 3) the rRNA platform is folded, and mitoribosomal proteins uL16m, bL36m, and mL88 are recruited, 4) MRM:mt-LAF15 is removed, uL11m and mL43 then adopt their mature conformations, 5) bL10m and bL12m are finally associated to form the functional L7/L12 stalk.

From the density we identified three additional proteins below the L7/L12 stalk: a homolog of the human mitoribosome assembly factor MALSU1, a LYR (leucine-tyrosine-arginine) motif containing protein L0R8F8, as well as an associated mt-ACP (mt-ACP1) (Figs 1, 2 and 4). In human and fungi, protein trans-acting factors in this region were shown to be involved in the last assembly stage of the mitoribosome, preventing association of the mtSSU (Brown et al 2017; Itoh et al 2020). In our structure, the module is further stabilized by mL85 to provide a steric hindrance, consistent with the previously suggested mechanism.

### Assembly of the CP is linked to the subunit interface and L1 maturation via mt-ACP

The most prominent architectural feature of the pre-mtLSU complex is the absence of the CP. It is a universal ribosomal element that defines the shape of the LSU and forms bridges with the SSU head. In mitoribosomes, the CP is particularly protein-rich (Amunts et 2014; Greber et al 2014; Amunts et 2015; Greber et al 2015; Ramrath et al 2018; Waltz et al 2020; Tobiasson and Amunts 2020). The CP mitochondria-specific proteins were acquired in the early evolution of the mitoribosome and therefore are expected to be conserved (Petrov et al 2019).

In the pre-mtLSU, all the CP mitoribosomal proteins are missing and the high molecular weight assembly factors mt-LAF4 (69 kDa) and mt-LAF14 (67 kDa) are present (Figs 1 and 4, Figure EV4). Factor mt-LAF4 binds on the solvent side of the CP covering the otherwise exposed rRNA, which only engages in limited base pairing. This assembly factor is annotated as a putative TruD family pseudouridine synthase. However, in our structure, it displays a two-strand antiparallel β-sheet near the catalytic site inserting into the ribosomal core and interacting with four other mitoribosomal proteins (Fig 8A). Due to this disruption of the active site the catalytic activity of mt-LAF4 is likely lost. Factor mt-LAF14 is an exclusively helical protein, comprised of at least 29 helices. It binds on top of the rRNA, providing an additional protective protein cap (Figs 1 and 8A).

**Figure 8.**
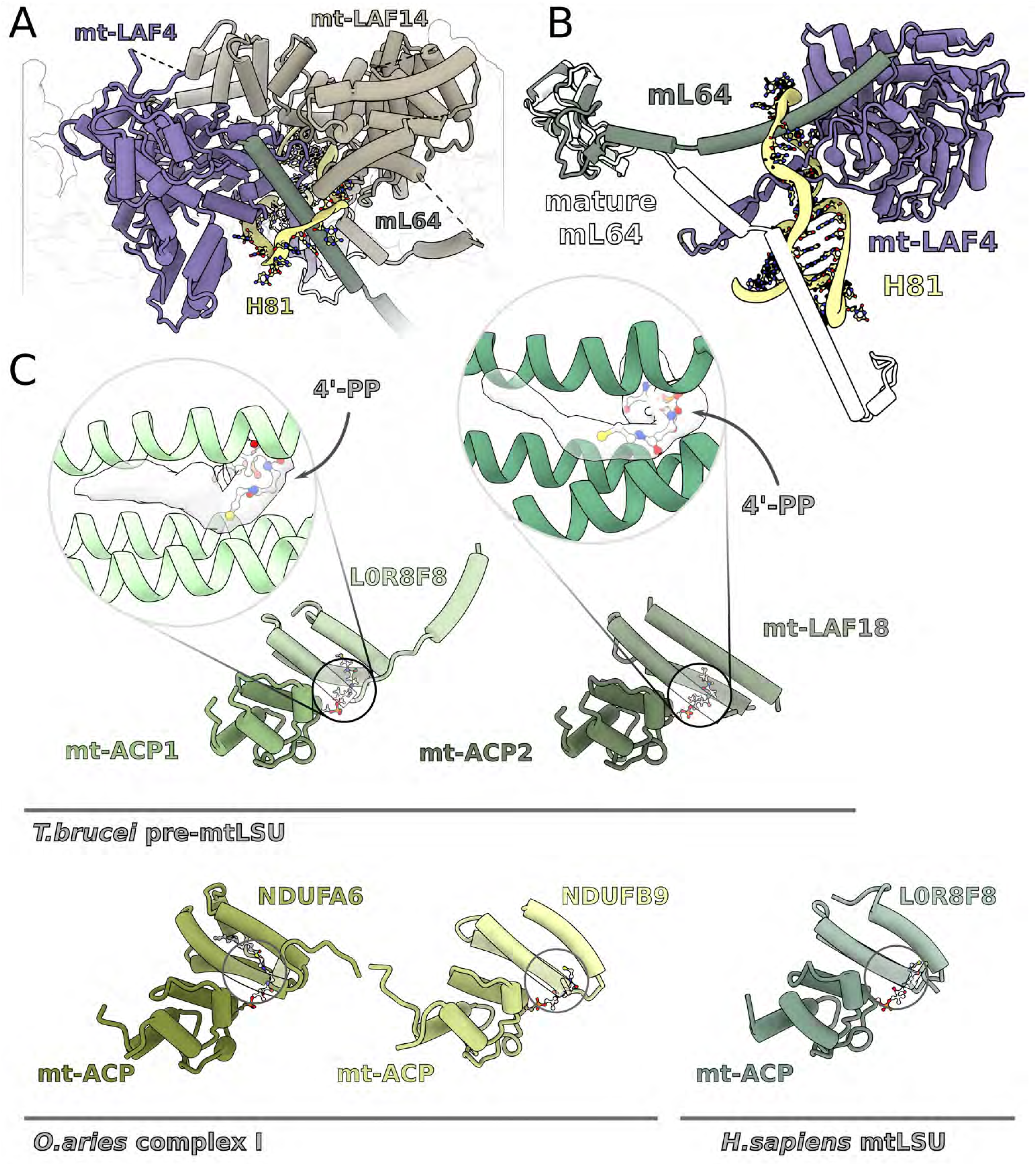
The CP assembly intermediate. (**A**) Factors mt-LAF4 and mt-LAF14 form the CP in the pre-mtLSU. (**B**) mt-LAF4 and mL64 elements are inserted through the rRNA loop corresponding to H81. The conformational change of mL64 from pre-mtLSU to mature mtLSU (white) is indicated. (**C**) Comparison between the mt-ACP1:L0R8F8 (left) and the CP mt-ACP2:mt-LAF18 region (right). The density (white) for acylated 4′-PP is indicated. Bottom panel, comparison with mt-ACP and associated LYR-motif proteins from complex I (PDB ID 5LNK) and human mitoribosome (PDB ID 5OOM) shows the canonical interactions.

Two of the mt-LAF14 helices interface with a four-helix bundle, which we identified as mt-ACP (mt-ACP2) with an average local resolution of 3.5 Å (Fig 2). Since mt-ACP proteins are known to form interactions with Leu-Tyr-Arg (LYR)-motif proteins, we compared the mt-ACP1:L0R8F8 module from the subunit interface with the CP mt-ACP2 region (Fig 8C). The helices of the LYR-motif protein L0R8F8 aligned well with a density corresponding to three helices associated with the mt-ACP2. The interactions in both cases are mediated by the 4′-phosphopantetheine (4′-PP) modification of mt-ACP, resembling the canonical interactions between mt-ACP and the LYR-motif proteins. The 4′-PP appears to be acylated as indicated by the density however the exact length of the acyl chain cannot be determined (Fig 8C).

The presence of the 4′-PP modification, previous structural data (Zhu et al 2015; Fiedorczuk et al 2016; Brown et al 2017), and the overall shape of the associated density suggest that the interacting partner of mtACP2 is a protein from the LYR-motif family containing at least three helices. Therefore, we searched our current and previously published (Zikova et al 2008) mass spectrometry data using ScanProsite (de Castro et al 2006). The hits were subjected to secondary structure prediction and fitting to the density map. Our analysis singled out the protein Tb927.9.14050 (UniProt ID Q38D50), which we named consistently with the adopted nomenclature, mt-LAF18. The local resolution of 3.5–4.0 Å in this region (Fig 2) allowed for 164 N-terminal residues to be built (Table EV2), which includes the three helices associated with the mt-ACP2 in proximity to the L1 stalk and two helices interacting with mt-EngA.

The importance of the mt-ACP2:mt-LAF18 protein module in our structure is of twofold. First, it directly binds the L1 stalk and locks it in an inward facing nonfunctional conformation (Fig 1). Second, it is involved in mt-EngA stabilization by forming a U-shaped continuum from mt-LAF14 on the solvent side of the CP to the subunit interface (Figs 1 and 4). Therefore, mt-ACP2 contributes to the protein network connecting between the various functional sites. In the pre-mtSSU, mt-ACP was also identified as one of the assembly factors (Saurer et al 2019). In addition, ACPs in mitochondria act as subunits of the electron transport chain (Zhu et al 2015; Fiedorczuk et al 2016) and Fe-S cluster assembly complexes (Van Vranken et al 2016). This further supports the proposed concept that mt-ACPs could be signaling molecules in an intramitochondrial metabolic state sensing circuit (Masud et al 2019).

At the CP, the assembly factors cooperatively bind unwound rRNA nucleotides U934-953 (H81) Figs 8A, EV4 and Appendix S6). Remarkably, the rRNA forms a loop 25 Å in diameter that encircles the mt-LAF4 β-sheet and mL64 C-terminal helix, both inserted from the opposite directions (Fig 8B). The conserved helix of mL64 is shifted in our pre-mtLSU structure ~30 Å from the final location on the mature LSU, where it aligns the E-site. Interestingly, this is also one of the most conserved mitoribosomal proteins (Petrov et al 2019). To switch to the conserved and mature position, the extended C-tail of mL64 has to liberate from the rRNA loop and then undergo a large conformational shift towards the L1 stalk. Subsequently, the C-tail is inserted to its mature position, where it contacts CP components absent from the assembly intermediate. Since the L1 stalk is also shifted, the maturation towards a mature mtLSU is likely to occur in a concerted manner upon the release of the mt-ACP2:mt-LAF18 module.

## Discussion

Our cryo-EM structure reveals how the assembly factors collectively bind to the mtLSU during biogenesis. High molecular weight assembly factors shield the rRNA and form a network that spans over 180 Å, which connects the subunit interface with the maturation of the L7/L12 stalk, and the assembly of the CP and the L1 stalk. The tight binding of the mt-ACP2 with its partner proteins, one from the CP and the other from the L1 stalk, emphasizes a coordinating role. Thus, the PTC is unfolded, the L1 is anchored in its inactive conformation, and the mitoribosomal proteins responsible for the binding of tRNA and translational factors cannot associate due to the presence of the assembly factors at the CP and L7/L12 stalk. In addition, the exit tunnel is blocked. In this regard, the present study is in agreement with the recently published work on *Leishmania* pre-mtLSU (Soufari et al 2020), which suggested that mL67, mL71, mL77, mL78, and mL81 represent assembly factors. The N-terminus of mL71 fills the exit tunnel, and its basic residues form electrostatic interactions with the rRNA that anchor the protein moiety. For a nascent polypeptide to emerge from the mtLSU, a continuous pathway needs to be formed from the tunnel entrance to the mitoribosomal surface, therefore mL71 N-terminus has to be removed.

Together, our pre-mtLSU structure contains 22 assembly factors, several of which could also be identified in the human mitoribosome assembly pathway, including GTPBP7, MRM, RPUSD4, MALSU1, L0R8F8, mt-ACP1, and a DEAD-box RNA helicase. This allowed us to suggest a model that underpins the organization of the equivalent assembly factors in the human pre-mtLSU (Fig 9). Functionally, these assembly factors can be divided into three categories: 1) GTPBP7 and DEAD-box helicase that potentially retained their functional role of facilitating rRNA folding; 2) MRM and RPUSD4, which lost their enzymatic functions, but retained the structural role of scaffolding the assembly process; 3) MALSU1, L0R8F8, and mt-ACP1 that form a conserved module preventing premature subunit association.

**Figure 9.**
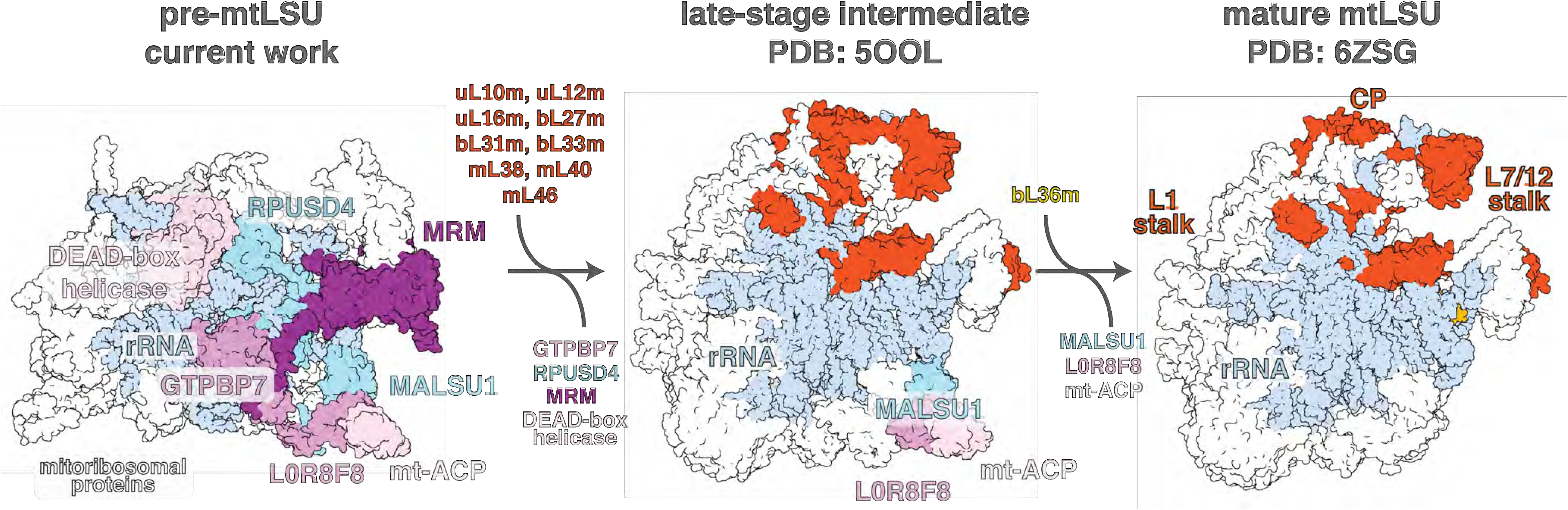
Schematic representation of the assembly pathway of human mtLSU. Left, the conserved assembly factors identified in this study that are also present in human are shown in complex with the pre-mtLSU. Middle, previously reported late assembly intermediate of the human mitoribosome (PDB 5OOL, Brown et al 2017) with assembled elements (relative to pre-mtLSU) shown in red. Right, mature mtLSU with fully assembled tRNA binding sites A, P, and E (PDB 6ZSG, Aibara et al 2020).

GTPBP7 is an essential mitoribosomal assembly factor that acts at an early assembly stage in yeast (Barrientos et al 2003; Kim and Barrientos, 2018), and also can associate with a mature LSU (Zeng et al 2018). Our analysis confirms that the residues in the nucleotide binding pocket are conserved in the GTPase domains (G1, G4), as well as in the P-loop and Switch II regions, and the nucleotide fits its pocket in our structure. DEAD-box helicase is also likely to act on an early assembly stage, as it is buried in the core of the pre-mtLSU. The DEAD-box motif is conserved, its conformation correspond to the RNA-binding state (Theissen et al 2008). This is consistent with the recently published structure of the pre-mtLSU (Jaskolowski et al 2020).

The binding of the GTPBP7 and DEAD-box helicase is stabilized by co-localized factors, including MRM and RPUSD4. In yeast, a single amino acid substitution in the SAM pocket of MRM1 abolishes its methyltransferase activity, but does not alter the formation of fully functional mitoribosomes (Lövgren and Wikström 2001), while deletion of MRM1 leads to a defective assembly (Sirum-Connolly and Mason 1993). *RPUSD4* is an essential gene in human cells, and it is a component of mitochondrial RNA granules (Zaganelli et al 2017). Our study points to structural roles of MRM and RPUSD4 in the assembly pathway of the mitoribosomes. MRM can also act as a docking site for the catalytically active methyltransferases MRM2/3 (Jaskolowski et al 2020), involved in 2′-*O*-ribose methylation of a nucleotide in the H92 loop (Rorbach et al 2014). The preservation of the deactivated factors is likely due to the evolutionary conservation of the sequential assembly (Fig 9), where RPUSD4 also forms a platform for DEAD-box helicase, as well as further stabilizes its expanded helicase 2 domain upon ATP hydrolysis. This mechanism is analogous to the evolutionary preservation of the autonomous 5S rRNA in bacterial ribosomes due to its role in assembly of the LSU where it guides the biogenesis pathway (Huang et al 2020).

In conclusion, our findings provide new insights into the conserved mtLSU biogenesis process. Protein extensions of the assembly factors and additionally incorporated protein linkers stabilize the key assembly factors of the mtLSU in the functional sites. Some of the factors, such as MRM and RPUSD4 lost their original function, and serve as structural mediators for the binding of the functionally active and conserved GTPBP7 and DEAD-box helicase. Therefore, the data also provides insight into the assembly of the human mitoribosome, where corresponding assembly intermediates are less stable. This showcases how the structural approach of studying stabilized intermediates is instrumental for understanding dynamic macromolecular processes that can be extrapolated to human homologs.

---

**Figure EV1.**
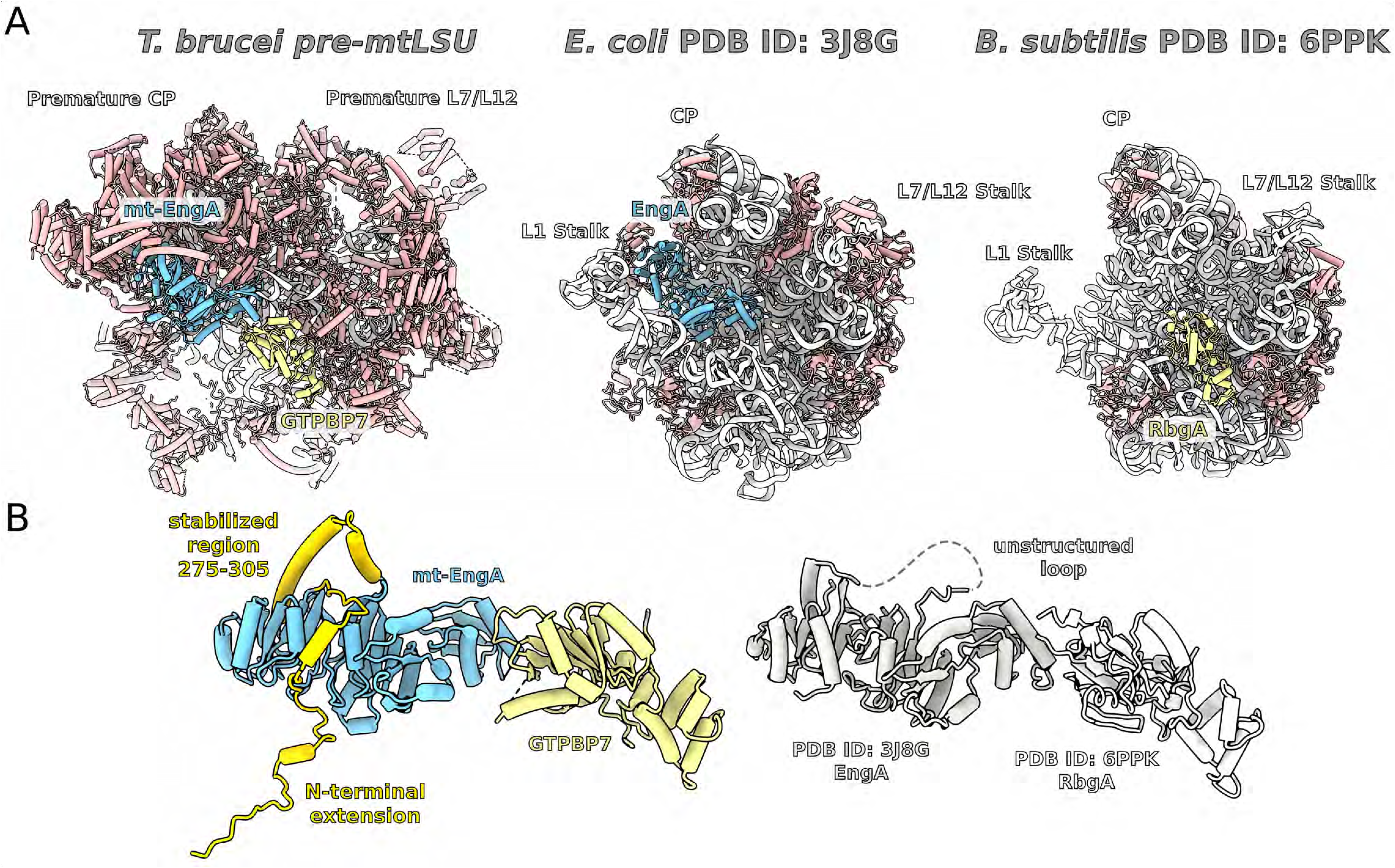
The binding of GTPBP7 and mt-EngA at the mtLSU interface. (**A**) Comparison between pre-mtLSU and bacterial counterparts *E. coli* 50S:EngA (PDB ID 3J8G) and *B. subtilis* 45S:RbgA (PDB ID 6PPK) shows nearly identical positions of the factors on their ribosomal complexes. (**B**) Comparison between GTPBP7:mt-EngA module from the pre-mtLSU and super-imposed bacterial counterparts combined from the two structures from (A) shows nearly identical conformations. The N-terminal extension of mt-EngA (dark yellow) is buried in the mitoribosomal core and stabilizes the binding, as well as the 275-305 region (dark yellow).

**Figure EV2.**
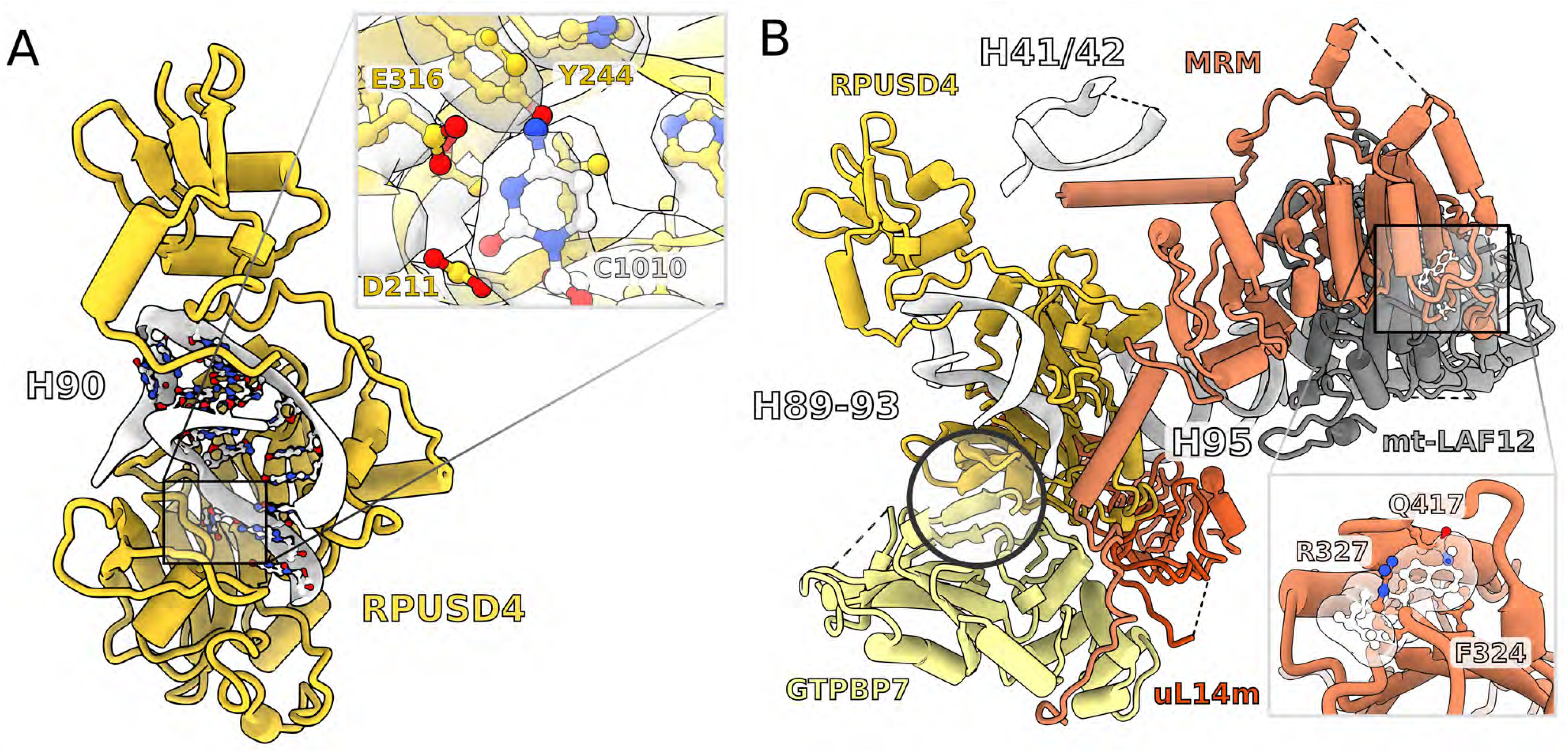
(**A**) The active site of RPUSD4 (yellow) is occupied by cytidine 1010. (**B**) The factor RPUSD4 (yellow) binds GTPBP7 (pale-yellow) via a shared β-sheet (circled). The methyl-transferase site of MRM does not allow for the binding of S-adenosyl methionine cofactor (white sticks and surface) due to clashes with the protein residues (inset).

**Figure EV3.**
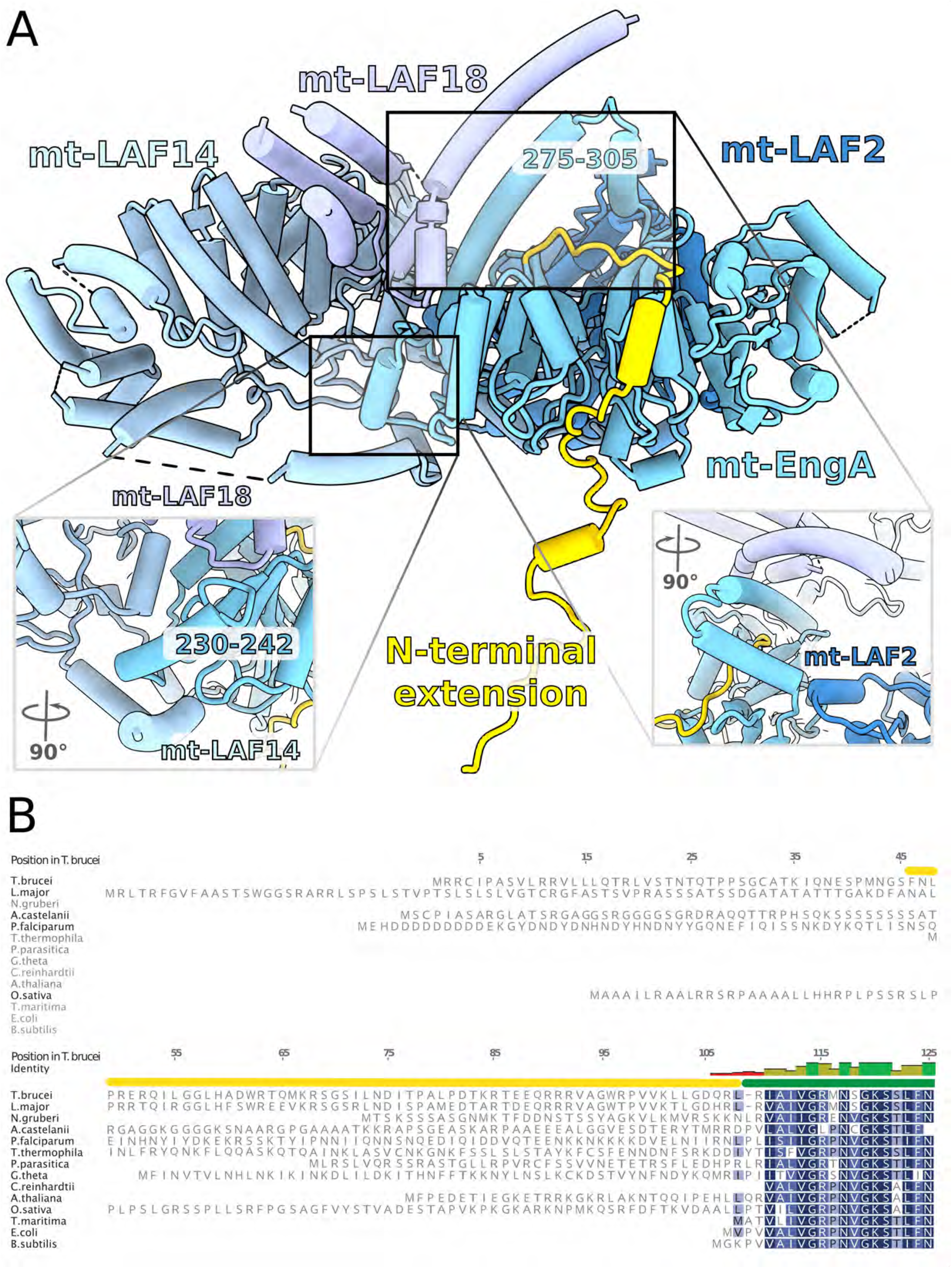
(**A**) N-terminal extension (yellow) of mt-EngA stabilizes helix-turn-helix (275-305), which forms interaction with mt-LAF2 on the other side (bottom right panel), and a helical bundle with mt-LAF18 that is in contact with mt-LAF14. (**B**) Sequence alignment of the N-terminus of mt-EngA shows presence of the extension in different organisms.

**Figure EV4.**
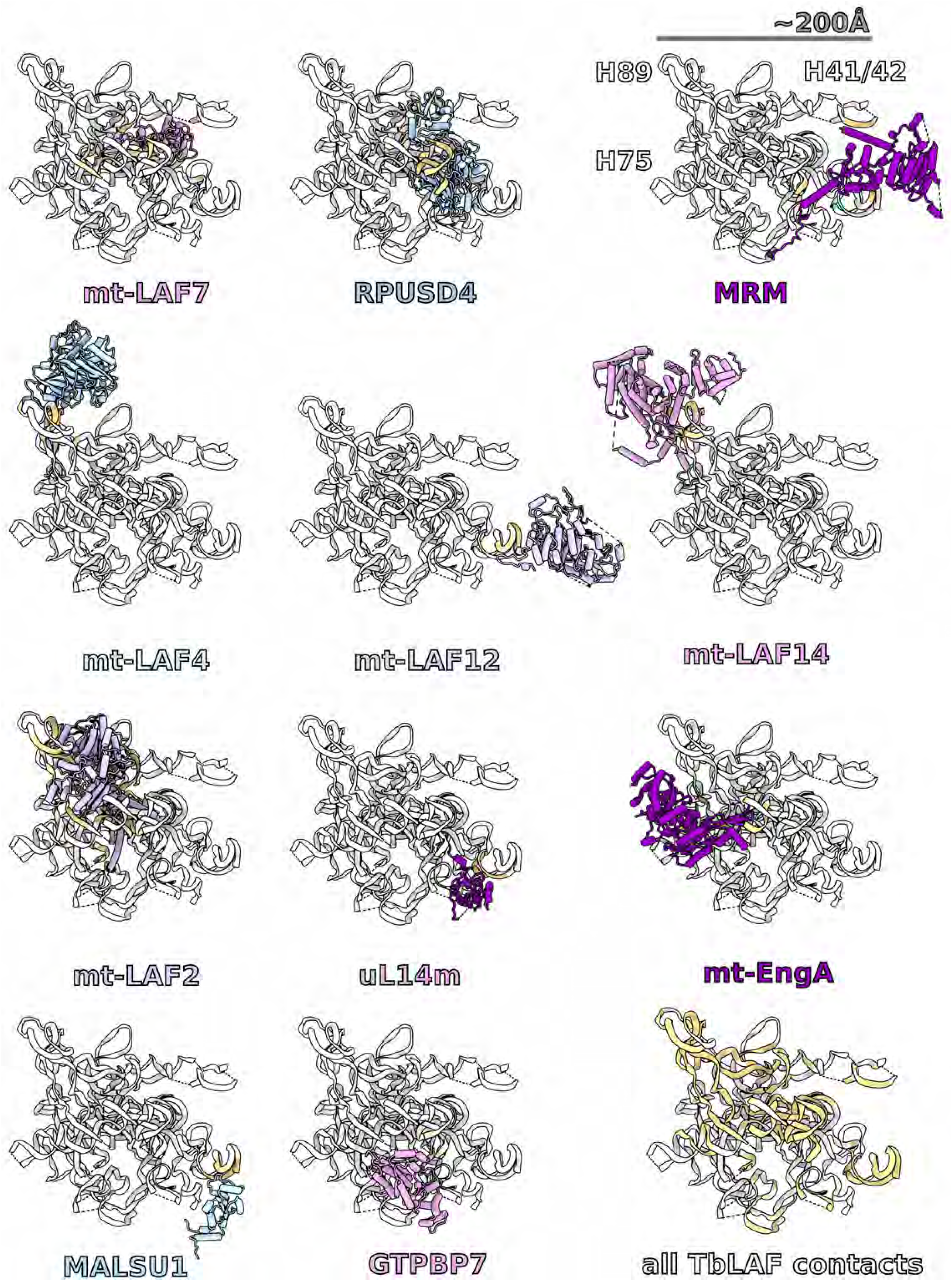
Binding of assembly factors to rRNA. For each panel, rRNA is shown with an individual protein characterized in the structure, which have not been reported in the mature LSU. Bottom right panel illustrates the total RNA that is involved in the interactions (yellow) with the assembly factors. Regions and nucleotides of respective rRNA domains are presented in Table S3.

**Figure EV5.**
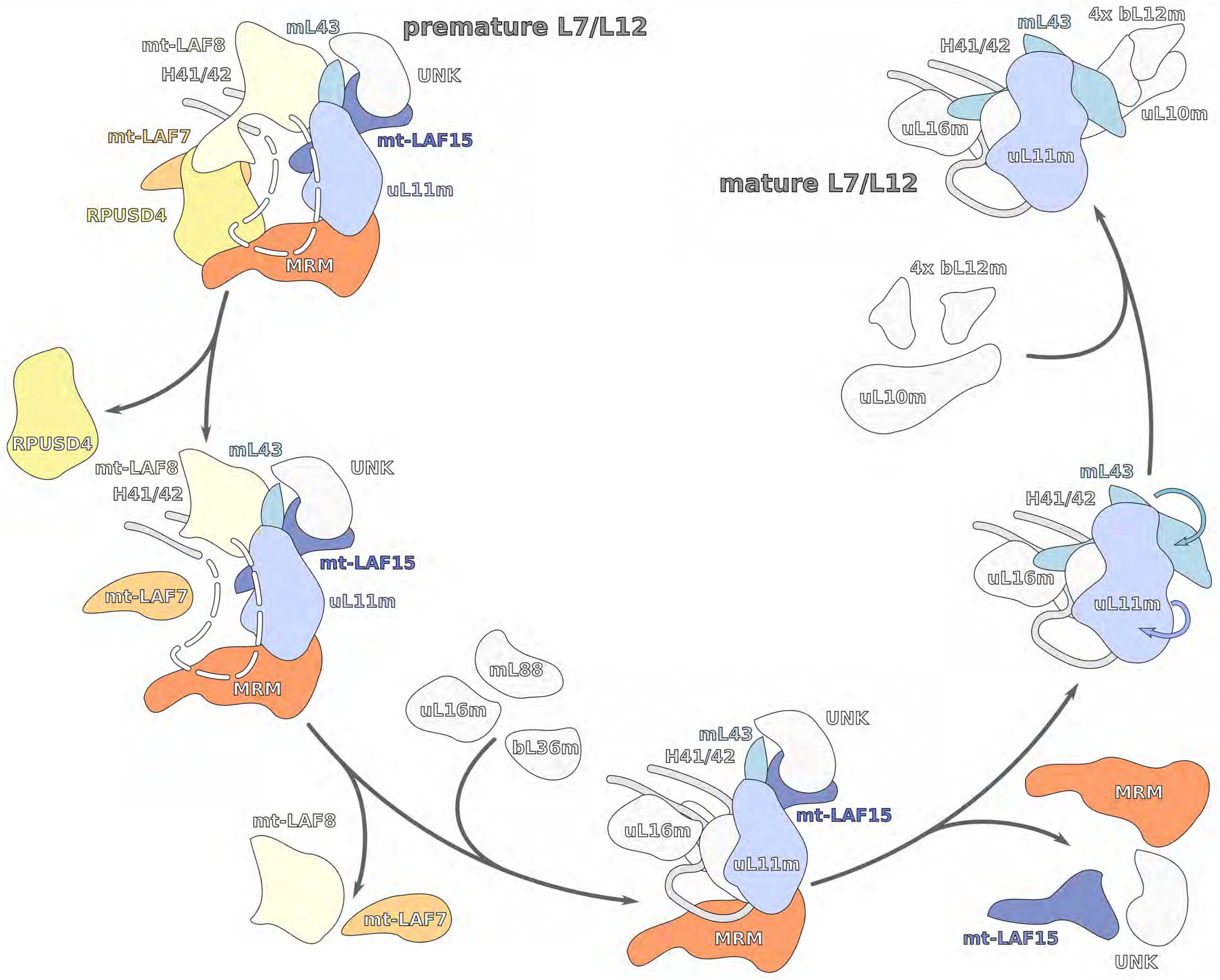
Proposed model for the L7/L12 stalk maturation. The series of steps starts with dismantling the assembly factors from the unfolded rRNA (white dashes) that triggers rRNA folding (grey line), binding of the mitoribosomal proteins (grey) and conformational changes (arrows).

## Materials and Methods

### Strains and growth conditions

*T. brucei* procyclic Lister strain 427 was grown in SDM-80 medium supplemented with 10% fetal bovine serum. Mitochondria were isolated as described earlier Schneider (2007). 1.5×10^11^ cells were harvested, washed in 20 mM sodium phosphate buffer pH 7.9 with 150 mM NaCl and 20 mM glucose, resuspended in 1 mM Tris-HCl pH 8.0, 1 mM EDTA, and disrupted by 10 strokes in 40 ml Dounce homogenizer. The hypotonic lysis was stopped by immediate addition of 1/6 volume of 1.75 M sucrose. Crude mitochondria were pelleted (15 min at 16000 xg, 4°C), resuspended in 20 mM Tris-HCl pH 8.0, 250 mM sucrose, 5 mM MgCl_2_, 0.3 mM CaCl_2_ and treated with 5 μg/ml DNase I for 60 min on ice. DNase I treatment was stopped by addition of one volume of the STE buffer (20 mM Tris-HCl pH 8.0, 250 mM sucrose, 2 mM EDTA) followed by centrifugation (15 min at 16000 xg, 4°C). The pellet was resuspended in 60% Percoll in STE and loaded on the bottom of six 10-35% Percoll gradient in STE in polycarbonate tubes for SW28 rotor (Beckman). Gradients were centrifuged for 1 hour at 24000 rpm, 4°C. The middle diffused phase containing mitochondrial vesicles (15-20 ml per tube) was collected, washed twice in the STE buffer, snap-frozen in liquid nitrogen and stored at −80°C.

### Purification of mitoribosomes

Mitochondria were purified further using a stepped sucrose gradient (60 %, 32 %, 23 %, 15%) in a low ionic strength buffer (50 mM HEPES/KOH pH 7.5, 5 mM MgOAc, 2 mM EDTA). A thick pellet at the 60-32% interface was collected and lysed by mixing with 5 volumes of detergent containing lysis buffer (25 mM HEPES/KOH pH 7.5, 100 mM KCl, 15 mM MgOAc, 1.7 % Triton X-100, 2 mM DTT, Complete-EDTA Free Protease Inhibitor). The lysate was centrifuged at 30,000 xg twice, retaining the supernatant after each spin. The supernatant was then subjected to differential PEG precipitation; PEG 10,000 was added to reach a concentration of 1.5 % (w/v) and incubated on ice for 10 mins, followed by a spin at 30,000 xg. The supernatant was transferred to a fresh tube, and PEG 10,000 was added to reach a concentration of 8 % (w/v) then incubated on ice for 10 mins, followed by a spin at 30,000 xg.

The pellet was then resuspended in 800 μl of lysis buffer and then layered onto a 34% sucrose cushion (25 mM HEPES/KOH pH 7.5, 100 mM KCl, 15 mM MgOAc, 1.0 % Triton X-100, 2 mM DTT, Complete-EDTA Free Protease Inhibitor) in a TLA120.2 centrifuge tube (0.4 ml of cushion per tube). Mitoribosomes were pelleted through the cushion by centrifugation at 231,550 xg for 45 min. Pelleted mitoribosomes were resuspended using a total of 100 μl of resuspension buffer (25 mM HEPES/KOH pH 7.5, 100 mM KCl, 15 mM MgOAc, 0.01 % β-DDM, 2 mM DTT). The resuspended mitoribosomes were then layered onto a continuous 15-30 % sucrose gradient and centrifuged in a TLS55 rotor for 120 min at 213,626 xg. The gradient was fractionated manually, and fractions containing mitoribosome as judged by the 260 nm absorbance were pooled and buffer exchanged in a centrifugal concentrator.

### Cryo-EM and model building

For cryo-EM analysis, 3 μL of the sample at a concentration of OD260 3.5, was applied onto a glow-discharged (20 mA for 30 seconds) holey-carbon grid (Quantifoil R2/2, copper, mesh 300) coated with continuous carbon (of ~3 nm thickness) and incubated for 30 seconds in a controlled environment of 100% humidity and 4 °C temperature. The grids were blotted for 3 seconds, followed by plunge-freezing in liquid ethane, using a Vitrobot MKIV (FEI/Thermofischer). The data was collected on a FEI Titan Krios (FEI/Thermofischer; Scilifelab, Stockholm, Sweden, and ESRF, Grenoble, France) transmission electron microscope operated at 300 keV, using C2 aperture of 70 μm; slit width of 20 eV on a GIF quantum energy filter (Gatan). A K2 Summit detector (Gatan) was used to collect images at a pixel size of 1.05 Å (magnification of 130,000X) with a dose of ~35 electrons/Å2 fractionated over 20 frames. A defocus range of 0.8 to 3.5 μm was applied.

19,158 micrographs (after bad images were removed based on real and reciprocal space features) were collected across 5 non-consecutive data acquisition sessions and processed together using RELION. 896,263 particles were picked using Warp and coordinates were imported into RELION for particle extraction at an initial binning factor of two. The particles were subjected to supervised 3D classification using references generated previously in a screening dataset, which was started based on the *T. brucei* cytosolic ribosome as an initial model. This crude separation classified the 207,788 particles as mtLSU-like, and the remaining as mature mtLSU-like, SSU-like or monosomes. This subset was subjected to auto-refinement separately to improve the angular assignments and then classified further using fine-angular searches with a solvent mask applied. From the mtLSU-like particles, 32,339 particles were retained as pre-mtLSU of good quality and the rest were discarded as non-particles. The retained pre-mtLSUs were then subjected to auto-refinement once more to improve the angles further, this time applying a solvent mask during the refinement procedure, and then the 3D reconstructions obtained were used as a reference for CTF refinement to improve the reconstruction. The final map was then estimated for local resolution using RELION and sharpened with a B-factor appropriate for the reconstruction as estimated automatically using the postprocessing procedure.

Model building was done using *Coot* 0.9 (Emsley et al 2010). First the model of the mature mtLSU (PDB ID:6HIX) was fitted to the density. Chains present in the pre-mtLSU were then individually fitted and locally refined. Additional chains were first identified using information from sidechain densities. First the map density, chemical environment and sidechain interactions were used to create probable sequences. Those sequences were then queried against *T. brucei* specific databases; potential hits were evaluated individually and finally assigned. Models were modeled de-novo. All models were refined iteratively using PHENIX (Liebschner et al 2019) realspace refinement and validated using MolProbity (Williams et al 2018). The data collection, model refinement and validation statistics are presented in Table S1. All figures were prepared either in Chimera (Pettersen et al 2004) or ChimeraX (Goddard et al 2018) with additional graphical elements created using Inkscape.

### Search for homologs of assembly factors and sequence alignments

Homologs of assembly factors found in our pre-mtLSU and identified by cryo-EM were searched in the NCBI protein database with Position-Specific Iterated BLAST (Altschul et al 1997) using sequences of individual factors from *T. brucei* as queries. The searches were targeted against selected genera. Sequence alignments were generated with the MUSCLE (Larkin et al 2007) algorithm in Geneious (Biomatters Ltd., New Zealand) and corrected manually.

## Data availability

The electron density map has been deposited in EMDB under accession code EMD-11845. The model has been deposited in PDB under accession code 7AOI. All data is available in the paper or Supplementary Information.

## Acknowledgements

We acknowledge the ESRF beamline CM01 for provision of beam time, and would like to thank especially Eaazhisai Kandiah and Michael Hons for the excellent continuous support. We also thank the SciLifeLab cryo-EM and mass spectrometry facilities, Nikhil Jain for comments. This work was supported by the Swedish Foundation for Strategic Research (FFL15:0325), Ragnar Söderberg Foundation (M44/16), European Research Council (ERC-2018-StG-805230), Knut and Alice Wallenberg Foundation (2018.0080), EMBO Young Investigator Program to A.A., and by and by Czech Science Foundation (18-17529S), ERD fund (CZ.02.1.01/0.0/0.0/16_019/0000759) to A.Z., and by Czech Science Foundation (20-04150Y) to O.G. The cryo-EM facility is funded by the Knut and Alice Wallenberg, Family Erling Persson, and Kempe foundations.

## Author contributions

Project conceptualization: OG, AZ, AA; Sample preparation for cryo-EM: OG, SA, AA; Data acquisition and processing: SA; Model building and validation: VT, OG, SA, RB; Structural data interpretation: VT, OG, AA; Manuscript writing and figure preparation: VT, OG, SA, RB, AZ, AA.

---

**Appendix Figure S1.**
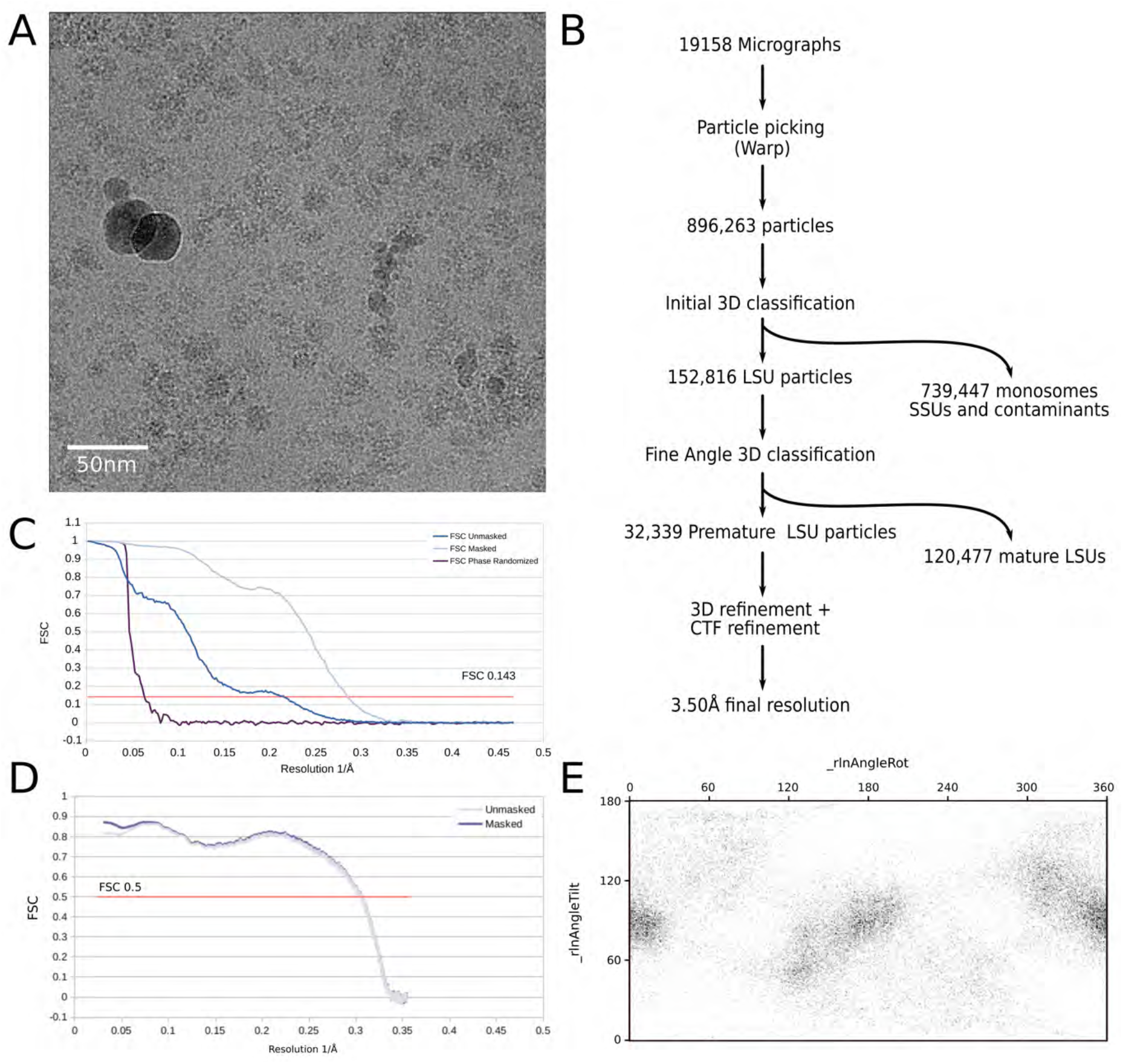
Cryo-EM data processing. (**A**) Representative micrograph. (**B**) Processing workflow. (**C**) Fourier shell correlation (FSC) curves. Resolution is estimated based on the 0.143 FSC cut-off criterion (red line).(**D**) Map to model FSC as calculated in PHENIX (Liebschner et al 2019). (**E**) Angular distribution plot for final reconstruction as calculated by RELION (Zivanov et al 2018).

**Appendix Figure S2.**
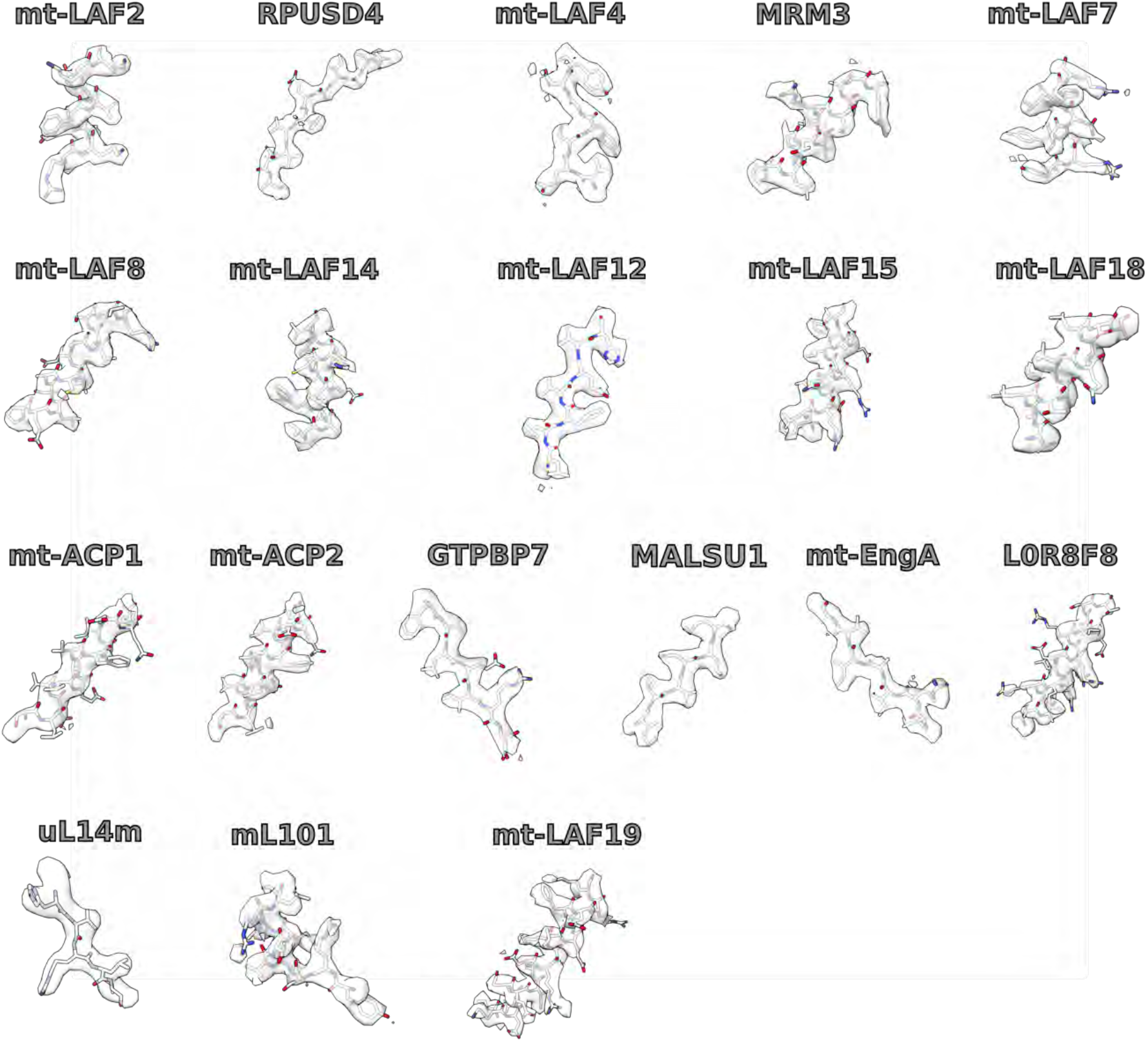
Examples of densities and models for individual assembly factors and newly identified proteins.

**Appendix Figure S3.**
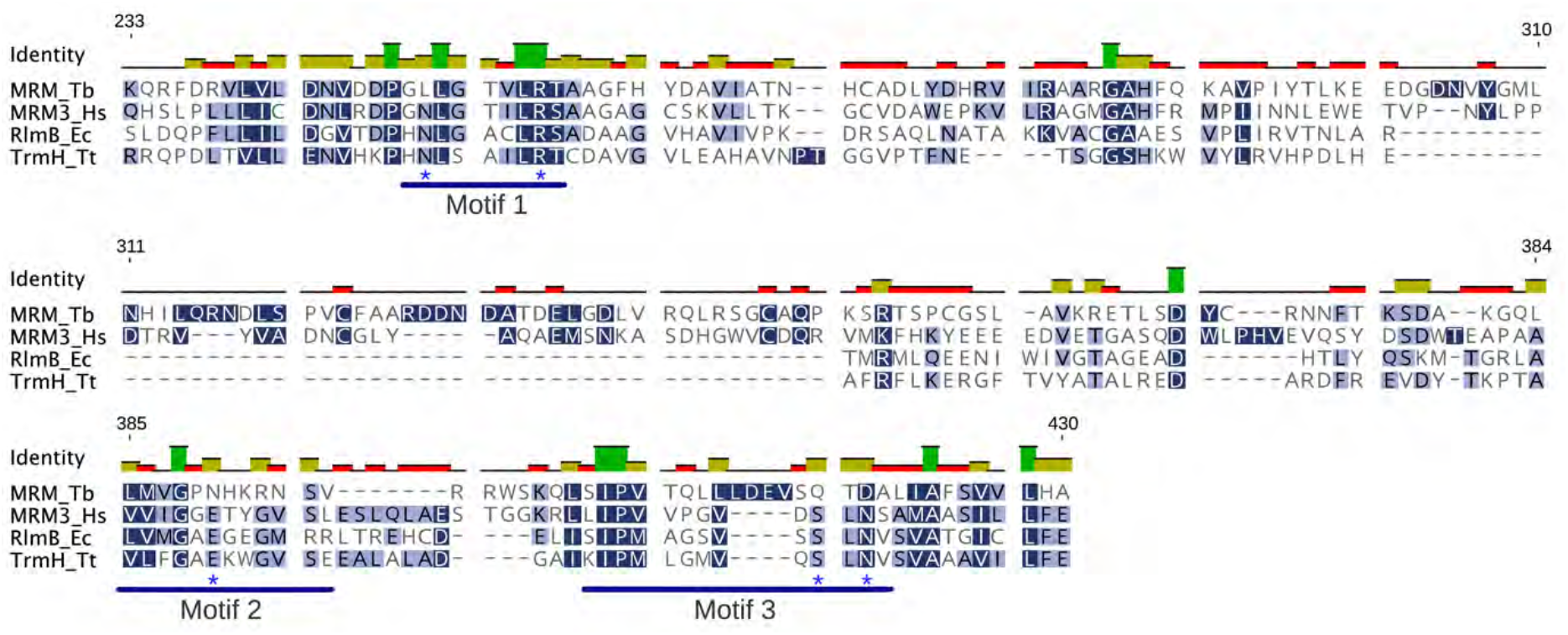
Sequence alignment of MRM homologs from representative bacterial and eukaryotic species (Hs *Homo sapiens*, Ec *E. coli*, Tt *Thermus thermophilus*. The asterisks mark residues important for catalysis.

**Appendix Figure S4.**
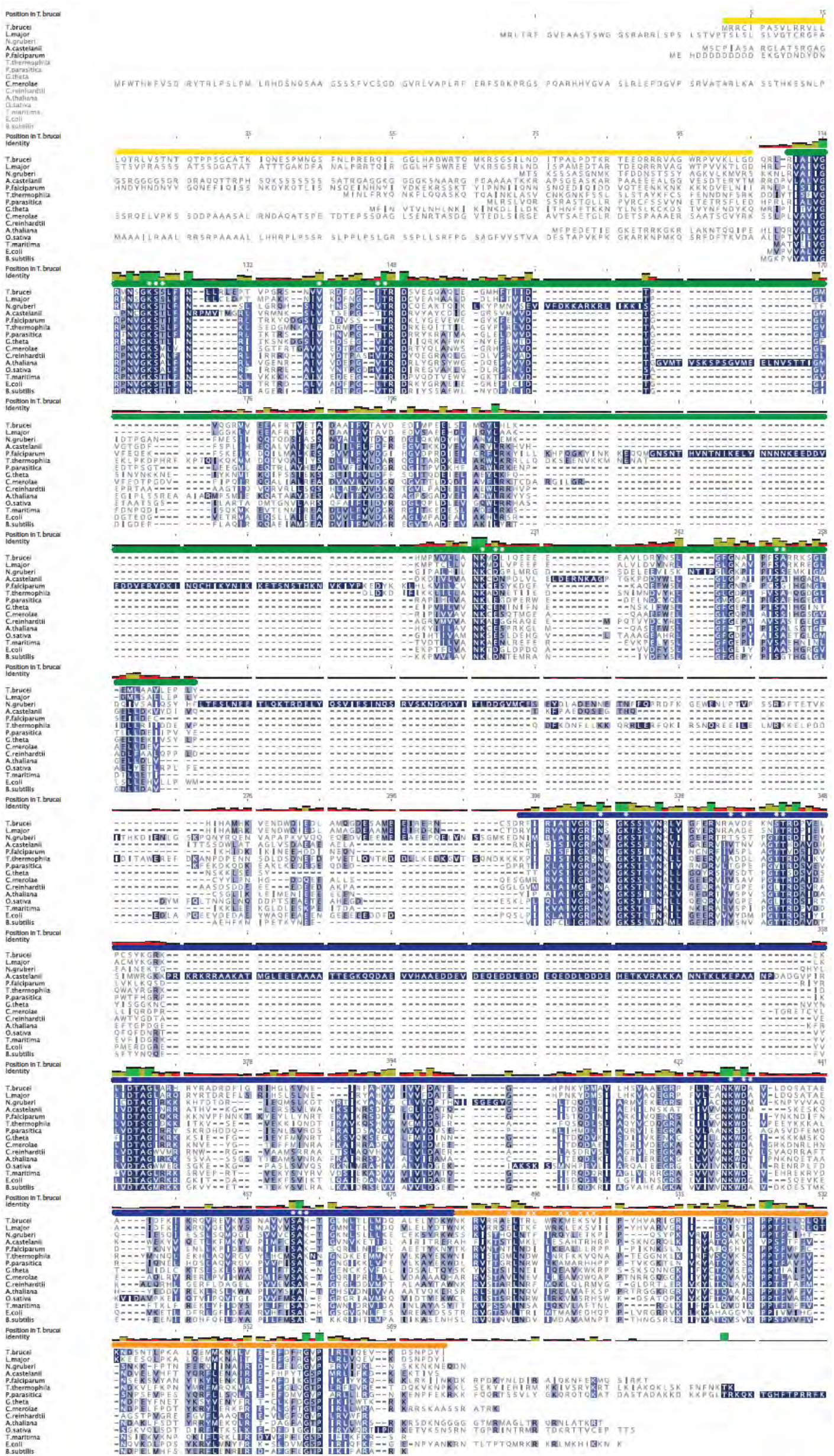
Sequence alignment of EngA homologs from representative bacterial and eukaryotic species. The yellow, green, blue and orange horizontal bars mark the N-terminal extension, GTPase domain (GD) 1, GD2, and the KH domain, respectively. The white asterisks and crosses mark residues in *T. brucei* mt-EngA that coordinate GTP and interact with GTPBP7, respectively. The green, yellow, and red vertical bars above the alignment correspond to 100%, <100% and ≥30%, and <30% identities at the respective position.

**Appendix Figure S5.**
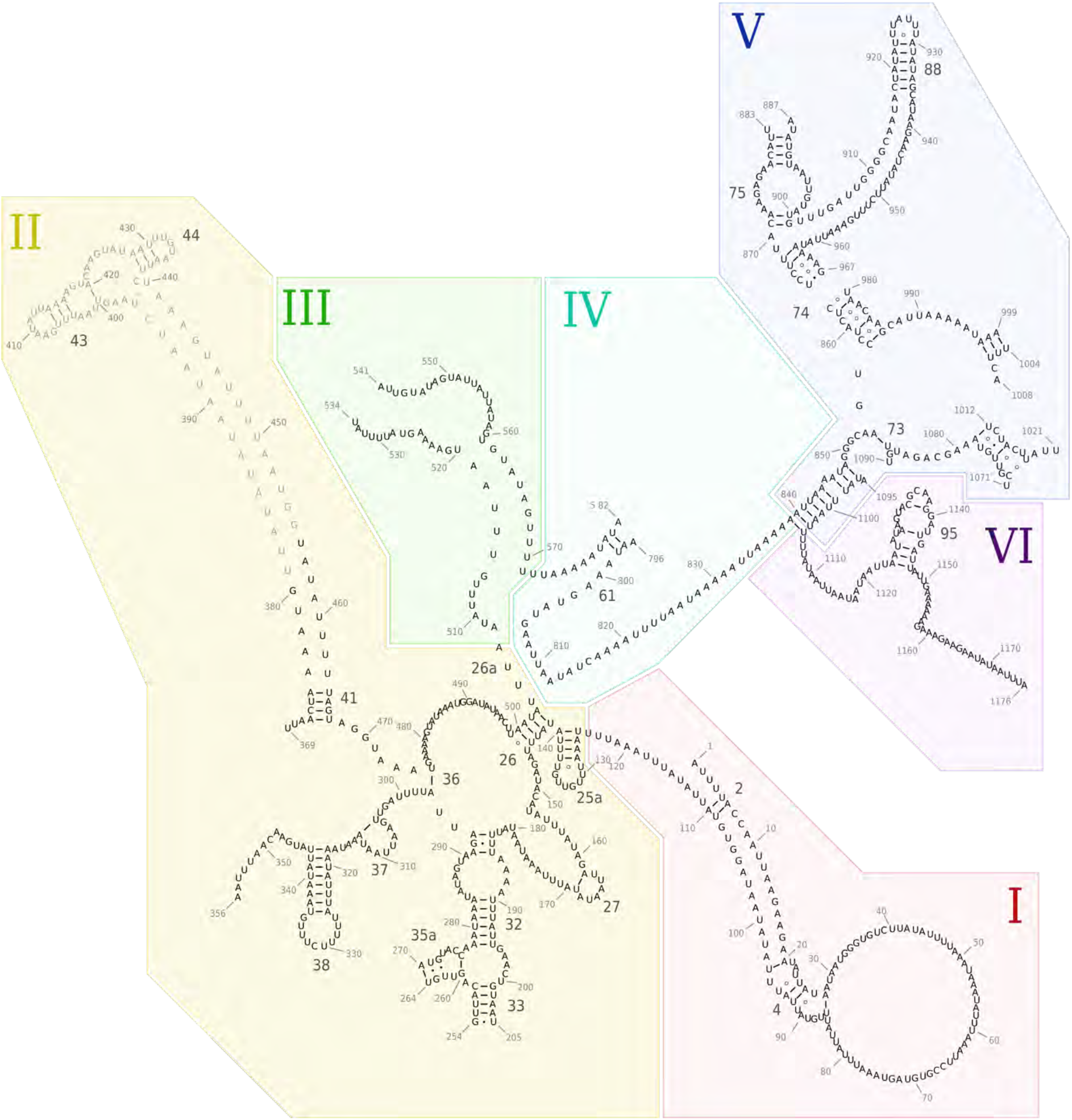
Secondary structure rRNA diagram derived from the model and colored by domain. Unmodeled sections that appear in the mature mtLSU are shown in grey. Domains in Roman numerals.

**Appendix Figure S6.**
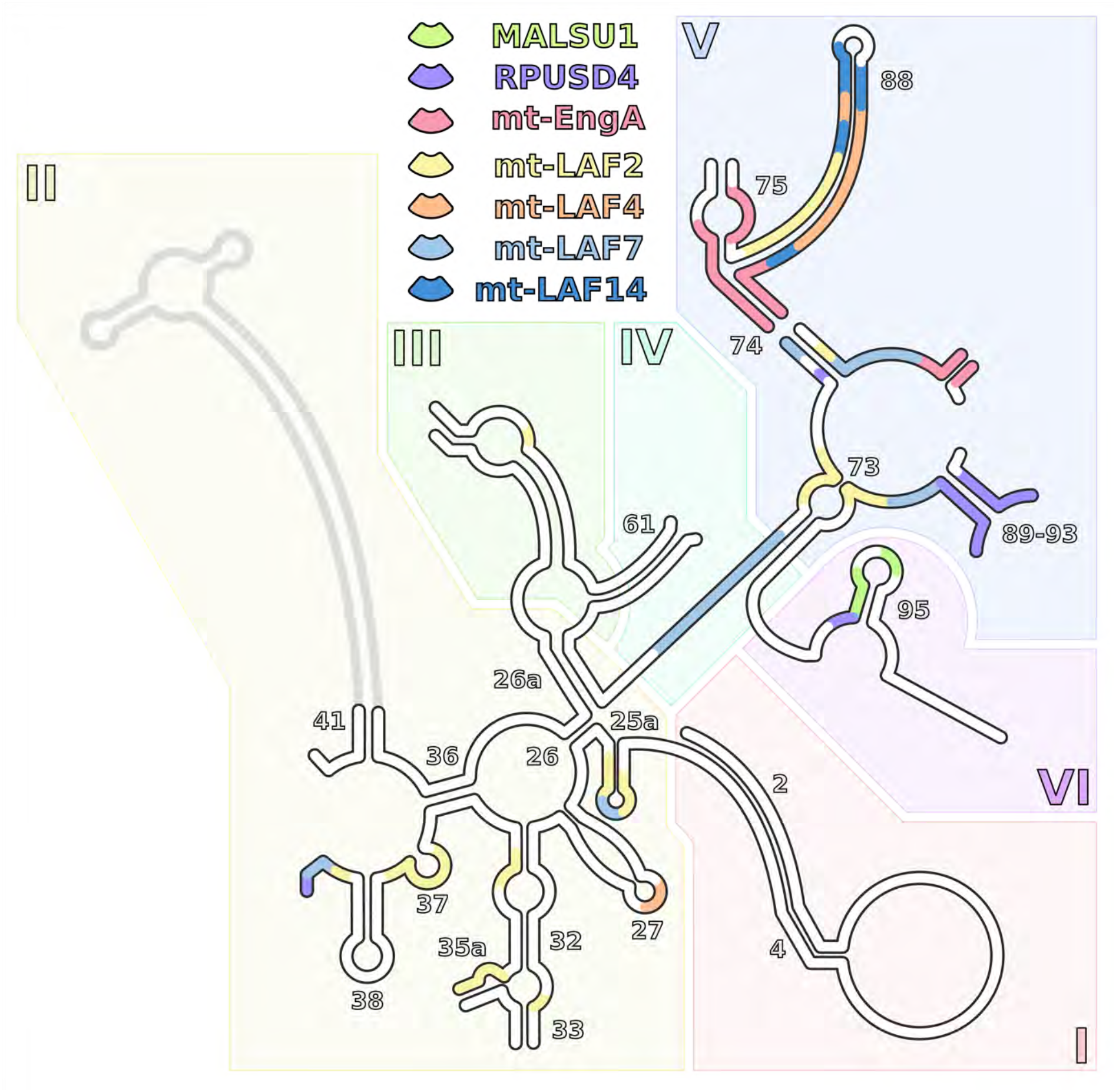
Schematic representation of assembly factors’ binding to rRNA mapped on the secondary structure diagram. The rRNA regions contacting individual assembly factors are represented by different colors. Bound regions of at least 3 nucleotides are shown. For regions where more than one factor is bound, only a factor with higher local binding is shown. Unbound rRNA is white, unmodeled rRNA is grey.

**Table S1.** Cryo-EM data collection, refinement and validation statistics.

|  |  |
| --- | --- |
| Consensus map |  |
| **Data collection and processing** |  |
| Magnification | 130000x |
| Voltage (kV) | 300 |
| Electron exposure (e-/Å<sup>2</sup>) | 35 |
| Defocus range (µm) | -0.8 ~ -3.5 |
| Pixel size (Å) | 1.05 |
| Symmetry imposed | C1 |
| Initial particle images (no.) | 896,263 |
| Final particle images (no.) | 32,339 |
| Map resolution (Å) | 3.50 |
| FSC threshold 0.143 |  |
| Map resolution range (Å) | 3.0~10 |
| **Refinement** |  |
| Map sharpening *B* factor (Å<sup>2</sup>) | -70 |
| Model composition |  |
| Non-hydrogen atoms | 146831 |
| Protein residues | 17415 |
| Ligands | 10 |
| *B* factors min/max/avg (Å<sup>2</sup>) |  |
| Protein | 17/172/68 |
| Nucleotide | 22/281/57 |
| Ligand | 35/194/61 |
| R.m.s. deviations |  |
| Bond lengths (Å) | 0.002 |
| Bond angles (°) | 0.46 |
| Validation |  |
| MolProbity score | 1.65 |
| Clashscore | 6.7 |
| Poor rotamers (%) | 0.35 |
| Ramachandran plot |  |
| Favored (%) | 95.6 |
| Allowed (%) | 4.10 |
| Disallowed (%) | 0.03 |

**Table S2.**
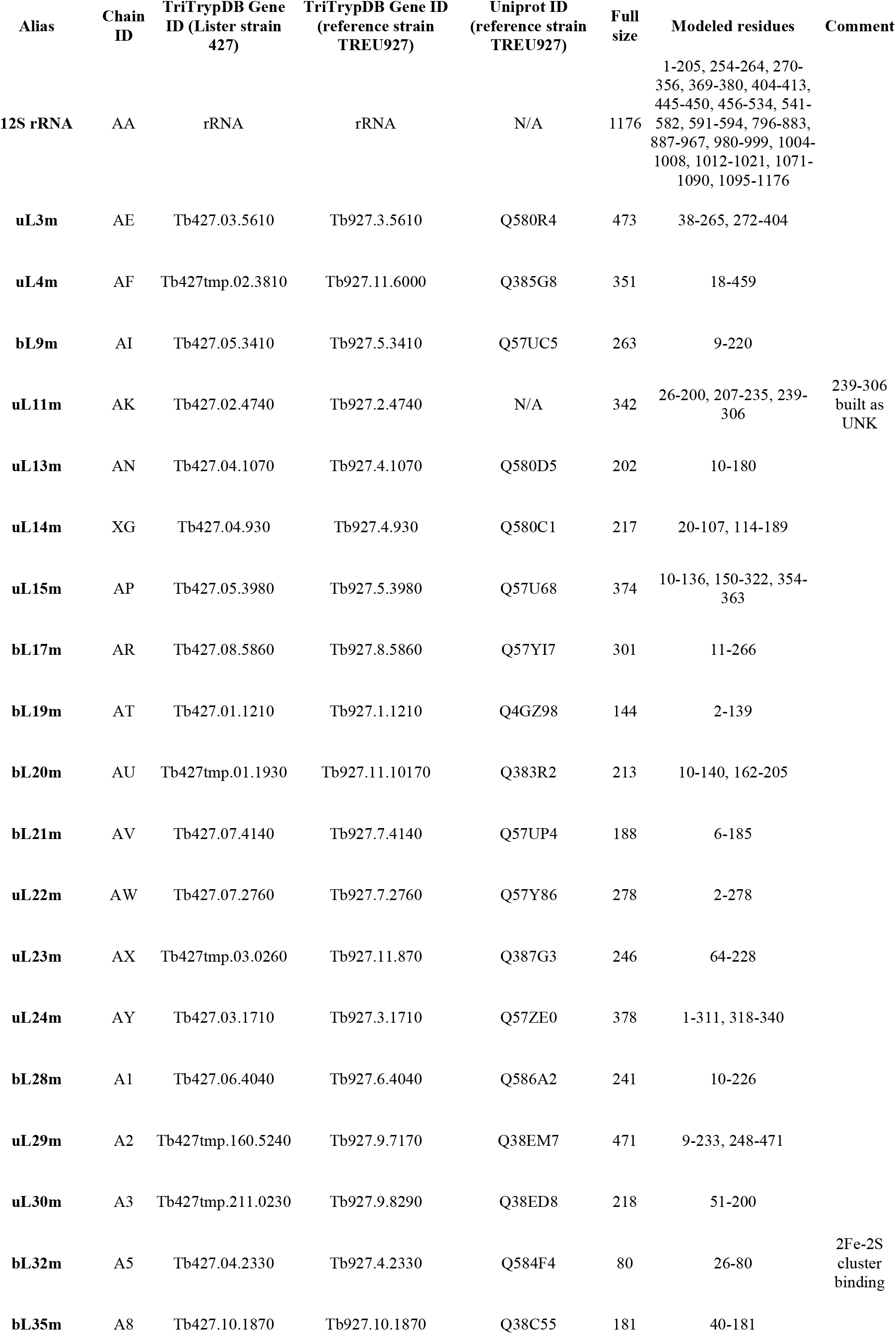
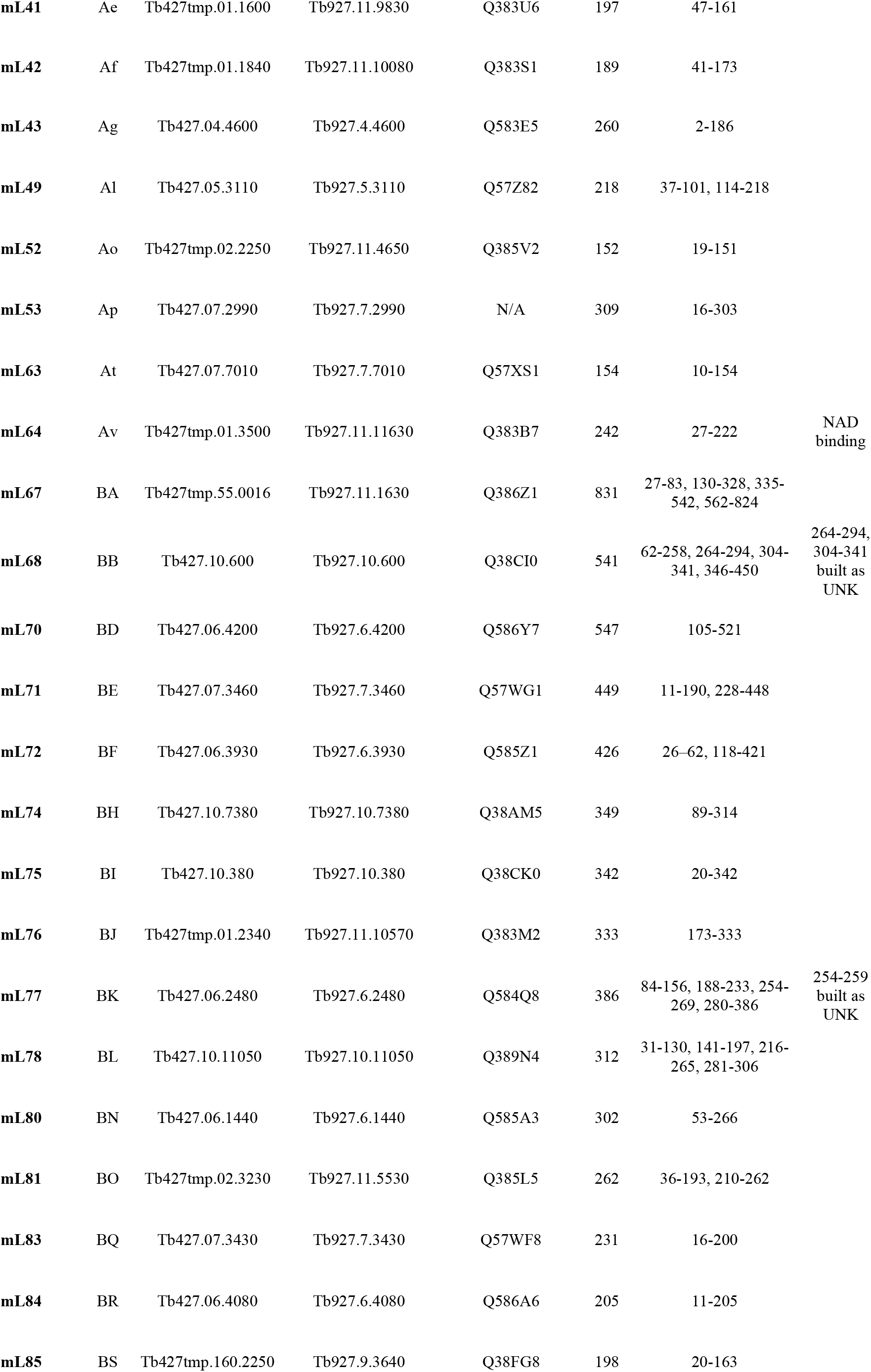
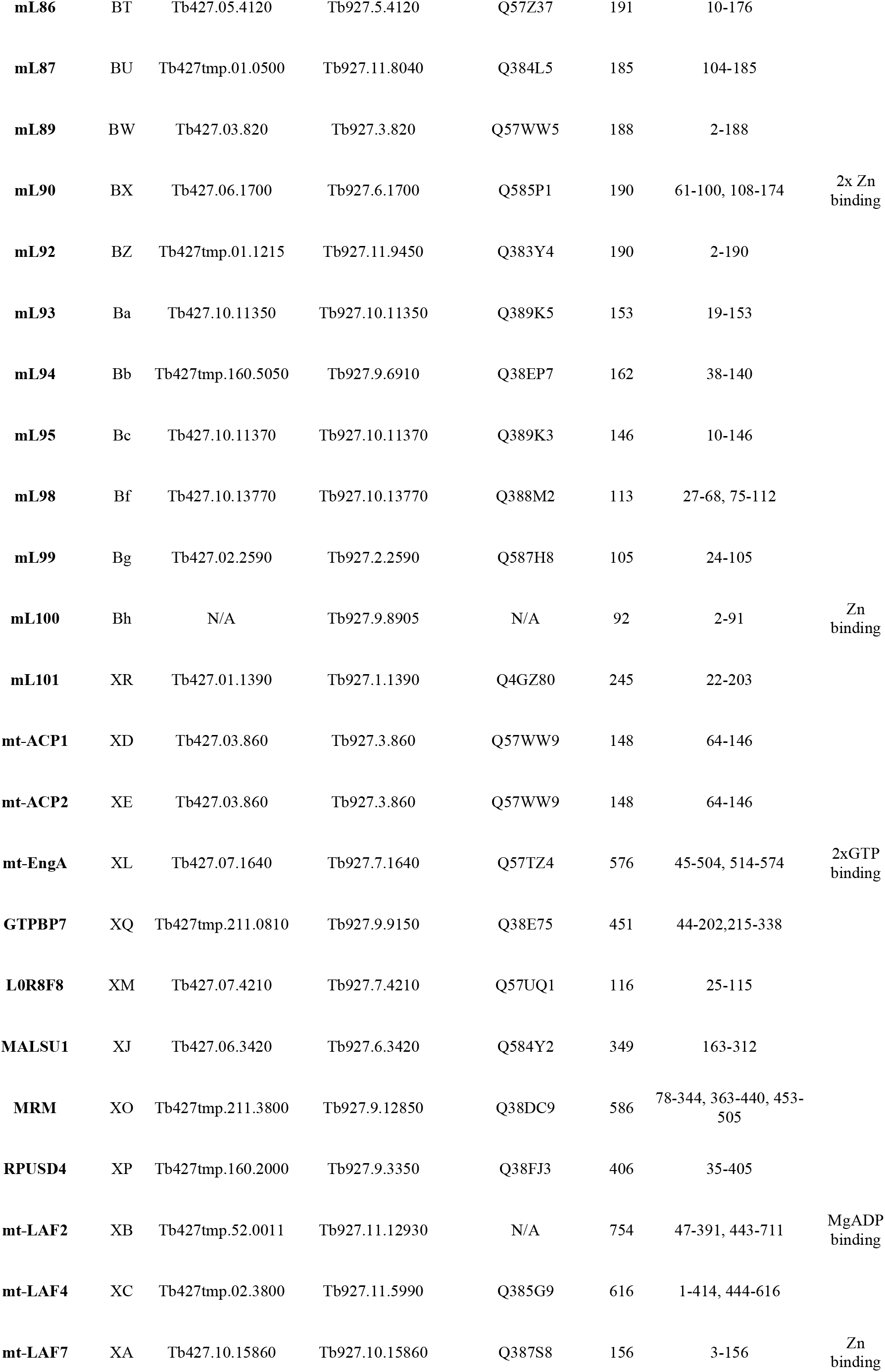
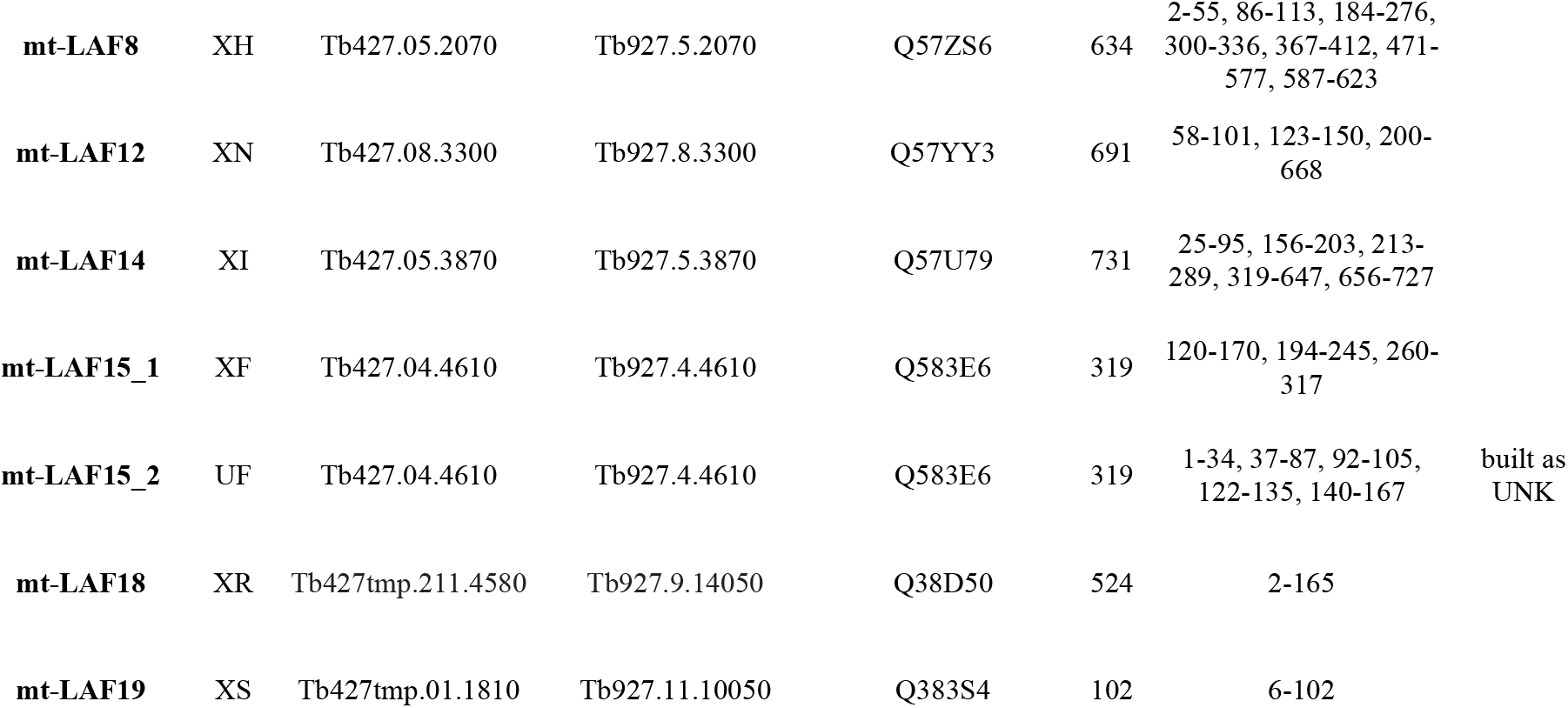
Summary of pre-mtLSU components.

**Table S3.** Contacts of assembly factors with rRNA. Regions and nucleotides of respective rRNA domains corresponding Fig EV4 and Appendix Fig S6.

| Assembly factor | Contacts with rRNA |  |
| --- | --- | --- |
|  | Region | Nucleotides |
| mt-EngA | H74, H75, H88, H90-91 | 868-874, 891-898, 958-967, 986-989 |
| GTPBP7 | H90-91 | 1013, 1071 |
| MALSU1 | H95 | 1126-1132, 1138-1141 |
| MRM | H43, H90, H95 | 409-413, 1012-1013, 1136-1137 |
| RPUSD4 | H39, H90-93 | 351-353, 857-859, 984-986, 1003, 1005-1013, 1071-1087, 1122-1124 |
| mt-LAF2 | H26, H32, H33, H35a, H37, H39, H51, H72, H73-75, H80-81, H93 | 127-139, 199-200, 270-276, 289-293, 306-316, 344-355, 497, 550, 555, 826-828, 847-848, 850-855, 860-862, 870, 903-922, 930-933, 947-950, 955-965, 975 - 984, 1085-1089 |
| mt-LAF4 | H13, H28, H37, H80-81, H88 | 65-66, 164-166, 313, 328, 908, 917-919, 935-939, 944-953 |
| mt-LAF7 | H39, H43, H72, H74, H88, H93 | 132-135 347-351, 404, 821-829, 832-835, 860-862, 904, 965-677, 977-984, 1079-1082, 1119 |
| mt-LAF8 | H41, H43 | 407, 448-450 |
| mt-LAF12 | H95 | 1131-1140 |
| mt-LAF14 | H80-81, H88 | 908-909, 915-916, 919-924, 930-940, 952-957, 961 |
| mt-LAF15 | H43 | 411-412, 446 |

## Notes

### Competing Interest Statement

The authors have declared no competing interest.

